# Sparse parallel independent component analysis and its application to identify linked genomic and gray matter alterations underlying working memory impairment in attention-deficit/hyperactivity disorder

**DOI:** 10.1101/2020.07.11.198622

**Authors:** Kuaikuai Duan, Jiayu Chen, Vince D. Calhoun, Wenhao Jiang, Kelly Rootes-Murdy, Gido Schoenmacker, Rogers F. Silva, Barbara Franke, Jan K. Buitelaar, Martine Hoogman, Jaap Oosterlaan, Pieter J Hoekstra, Dirk Heslenfeld, Catharina A Hartman, Emma Sprooten, Alejandro Arias-Vasquez, Jessica A. Turner, Jingyu Liu

## Abstract

Most psychiatric disorders are highly heritable and associated with altered brain structural and functional patterns. Data fusion analyses on brain imaging and genetics, one of which is parallel independent component analysis (pICA), enable the link of genomic factors to brain patterns. Due to the small to modest effect sizes of common genetic variants in psychiatric disorders, it is usually challenging to reliably separate disorder-related genetic factors from the rest of the genome with the typical size of clinical samples. To alleviate this problem, we propose sparse parallel independent component analysis (spICA) to leverage the sparsity of individual genomic sources. The sparsity is enforced by performing Hoyer projection on the estimated independent sources. Simulation results demonstrate that the proposed spICA yields improved detection of independent sources and imaging-genomic associations compared to pICA. We applied spICA to gray matter volume (GMV) and single nucleotide polymorphism (SNP) data of 341 unrelated adults, including 127 controls, 167 attention-deficit/hyperactivity disorder (ADHD) cases, and 47 unaffected siblings. We identified one SNP source significantly and positively associated with a GMV source in superior/middle frontal regions. This association was replicated with a smaller effect size in 317 adolescents from ADHD families, including 188 individuals with ADHD and 129 unaffected siblings. The association was found to be more significant in ADHD families than controls, and stronger in adults and older adolescents than younger ones. The identified GMV source in superior/middle frontal regions was not correlated with head motion parameters and its loadings (expression levels) were reduced in adolescent (but not adult) individuals with ADHD. This GMV source was associated with working memory deficits in both adult and adolescent individuals with ADHD. The identified SNP component highlights SNPs in genes encoding long non-coding RNAs and SNPs in genes MEF2C, CADM2, and CADPS2, which have known functions relevant for modulating neuronal substrates underlying high-level cognition in ADHD.

## 1. Introduction

Advancements in brain imaging and multi-omics techniques have significantly improved our understanding of psychiatric disorders, which consist of highly complex traits of clinical symptoms and associated genetic and neurobiological alterations. For many psychiatric disorders, including schizophrenia, bipolar disorder, and attention-deficit/hyperactivity disorder (ADHD), patients present widespread rather than focal brain alterations. These brain alterations often manifest as reduced gray matter volume (Chen et al., 2019b; Duan et al., 2018; Gupta et al., 2015; Hoogman et al., 2017; Hoogman et al., 2019; Jiang et al., 2020), altered functional connectivity (Chen et al., 2018; Fu et al., 2018), and low integrity of white matter tracts (White et al., 2011; Wu et al., 2015). In parallel, genome-wide association studies (GWAS) have revealed that genetic risk to most psychiatric disorders is shaped by many variants with small to modest effect sizes, known as polygenicity (Demontis et al., 2019; Grove et al., 2019; Ripke et al., 2014; Wray et al., 2018). Heterogeneous clinical symptoms are well recognized in most DSM-5 diagnostic categories. These findings lend support to dimensional frameworks to characterize psychiatric disorders in a multitude of dimensions where genetic, neurobiological and phenotypic factors are better integrated (Insel et al., 2010). Imaging genetic/genomic studies (Bogdan et al., 2017; Chen et al., 2019a; Liu and Calhoun, 2014; Mufford et al., 2017) play an important role in this framework, as they serve to delineate genetic underpinnings of neurobiological abnormalities linked to phenotypic manifestation. However, achieving this goal has proven to be a great challenge given the complexity of the diseases and modest effect sizes of individual genetic variants. Compared to univariate analysis methods (e.g., voxel based morphometry (Good et al., 2001) and GWAS (Bush and Moore, 2012)), multivariate data-driven fusion methods (Liu and Calhoun, 2014) pose a promising solution by aggregating related variables with small effect sizes into one network/source and simultaneously identifying brain imaging-genetic associations at the network level. Thus, multivariate fusion methods can boost the statistical power and facilitate the interpretation of findings.

Several data-driven multivariate fusion approaches have been proposed for imaging genetics, including sparse canonical correlation analysis (sCCA) (Chi et al., 2013; Du et al., 2016), sparse partial least squares (sPLS) (Le Floch et al., 2012), sparse reduced rank regression (sRRR) (Vounou et al., 2010) and parallel independent component analysis (pICA) (Liu et al., 2009; Pearlson et al., 2015). SCCA, sPLS and sRRR are extensions of CCA, PLS and RRR, respectively, which implement a sparsity constraint exploiting *l*_1_ norm to alleviate uncontrolled overfitting in the original methods to some extent. CCA and PLS maximize the correlation and covariance between two modalities, respectively. RRR minimizes the error of the multivariate regression by reducing the rank of the projection matrix mapping the input to the output. PICA differs from the aforementioned approaches in that it prioritizes extracting maximally independent components from individual modalities, while simultaneously optimizing the correlations between multimodal component expressions. The advantages and limitations of each method have been reviewed elsewhere (Liu and Calhoun, 2014). We note that sCCA, sPLS and sRRR are defined across modalities and thus limited to decomposing the same number of factors/components for all modalities, while pICA allows different numbers of components for different modalities. Moreover, when the intrinsic dimensionality of a modality is much larger than the sample size, such as with genomic single nucleotide polymorphism (SNP) data, the search space becomes so large that even methods such as sCCA, sPLS and sRRR are susceptible to overfitting. The very large dimensionality condition also makes it difficult for pICA to separate signals of interest (e.g., SNPs involved in a specific brain alteration) from the background genomic profile. Since the variance of genotypes carried by SNPs of interest is typically comparable to, or slightly higher than, that of background SNPs (i.e., SNPs modulating other or unknown biological processes), the independent components from pICA tend to assign similar or slightly higher weights for SNPs of interest compared to background SNPs (Duan et al., 2019). In this study, we propose a sparsity regulated pICA to mitigate issues with signalbackground-separation. Like pICA, the use of independence leverages higher-order statistics to overcome overfitting issues, while (nonlinear) sparsity regularization yields a crucial denoising effect for better signal detection.

Among sixteen sparsity measures that have been summarized in (Hurley and Rickard, 2009), we selected the Hoyer index as the most attractive, since it satisfies five out of six desirable axiomatic attributes of sparsity measures while also being differentiable and, thus, amenable to optimization. The Hoyer index is a robust extension of the ratio between *l*_1_ and *l*_2_ norms. Comparing to *l*_1_ norm, Hoyer index is scale invariant, which is well suited for ICA frameworks since the scale of decomposed components is arbitrary. Our previous work (Duan et al., 2019) has demonstrated that incorporating Hoyer index into infomax ICA improved the SNP component detection accuracy under various scenarios. Thus, we propose sparse parallel ICA (spICA) to incorporate robust sparsity optimization into the pICA framework, not only taking advantage of correlation optimization but also leveraging the desirable sparsity properties of the Hoyer index for nonlinear denoising. The effectiveness and the robustness of spICA are examined with simulated imaging and genetic data.

To examine the capability of the proposed spICA in real data, we apply spICA to structural brain imaging and SNP array data in ADHD cohorts. ADHD, a childhood-onset neuropsychiatric disorder, affects 5% to 7% children of school-age in the world (Polanczyk et al., 2007). In about 60% of children with ADHD, their symptoms persist into adulthood (Sibley et al., 2017). Besides the symptomatic persistence, working memory impairments (likely along with other cognitive impairments) may also persist from childhood (Alderson et al., 2017) to adulthood (Alderson et al., 2013; Mostert et al., 2015). Cumulative evidence from brain imaging studies supports that brain structural and functional differences indeed underlie overt symptoms and cognitive impairments (Chen et al., 2018; Duan et al., 2018; Fu et al., 2018; Goghari et al., 2014; Hoogman et al., 2019; Takeuchi et al., 2017). Our previous work (Duan et al., 2018) has identified that gray matter alterations in superior/middle/inferior frontal regions and cerebellum are related to working memory deficits or inattention symptoms in both children and adults with ADHD. Furthermore, ADHD is highly heritable, with an estimated heritability of 77–88% (Faraone and Larsson, 2019). A recent ADHD GW AS (Demontis et al., 2019) identified 12 independent loci showing significant genomic differences in cases vs. controls, with many promising loci contributing to ADHD susceptibility. However, each locus only accounted for a very small portion of the variance related to the disorder. In turn, the use of univariate methods for the identification of individual loci related to brain abnormalities in ADHD is largely hindered in current studies due to small sample sizes. Important work led by the Enhancing NeuroImaging Genetics through Meta-Analysis (ENIGMA) ADHD and schizophrenia consortiums (Hoogman et al., 2020; Turner, 2019) seeks to deal with the sample size problem through world-wide collaborations. Meanwhile, this work explores the equally important task of improving current multivariate methods, here pICA, via the introduction of a robust sparsity regularization for suppression of background information. We apply spICA to brain gray matter images and SNP array data, seeking to group modest effect size interactive genomic variants into one source while enhancing signals of interest that contribute to altered brain networks.

In this work, spICA is performed on structural magnetic resonance imaging (sMRI) and SNP array data of 341 unrelated adults aggregated from the NeuroIMAGE (von Rhein et al., 2015) and IMpACT-NL (Mostert et al., 2015; Onnink et al., 2014) projects. Specifically, three previously reported gray mater networks (Duan et al., 2018; Liu et al., 2020), which are consistently associated with working memory impairments and inattention in both adult and adolescent ADHD in this sample, are investigated together with SNPs comprehensively selected based on ADHD GWAS summary statistics (Demontis et al., 2019), to uncover genetic factors underlying brain alterations related to persistent ADHD in adults. A replication dataset consisting of sMRI and SNP data of 461 adolescents recruited by the NeuroIMAGE project (von Rhein et al., 2015) is employed to validate discovered sMRI-SNP pairs. Subsamples partitioned by age in the replication dataset are also tested to validate the association between SNP components and gray matter networks.

## 2. Materials and Methods

In this section, we first describe the proposed spICA algorithm, then introduce the simulation design, and subsequently detail the steps for real data analysis.

### 2.1. Sparse parallel independent component analysis (spICA)

The spICA proposed here is an extension of pICA (Liu et al., 2009), which imposes additional sparsity constraints on components/sources. PICA maximizes component independence for each modality in parallel (i.e., multiple ICAs) while simultaneously enhancing the intercorrelation between specific loadings across modalities. In spICA, the sparsity constraint is enforced at each iteration by applying the Hoyer projection (Duan et al., 2019; Hoyer, 2004) on the estimated components directly, which enhances signals of interest and effectively suppresses background information. This choice of implementation yields a net non-linear relation between components and the input data. Consequently, solutions otherwise unavailable from typical linear transformation solvers become attainable with spICA.

The cost function of the proposed spICA is shown in equation 1. SpICA maximizes the sum of differential entropies (H(**y**_1_) + H(**y**_2_)) and square of the intercorrelation between modalities (corr^2^(**a**_1_, **a**_2_)) with respect to unmixing matrices (**W**_1_, **W**_2_), subject to Hoyer index constraints on the components of both modalities (Hoyer(**s**_1_) ≥ q_1_ Hoyer(**s**_2_) ≥ q_2_).

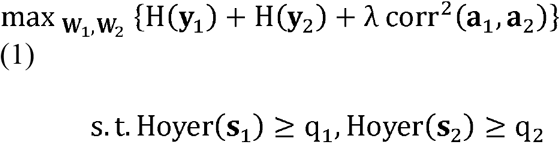

where

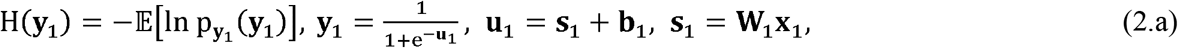

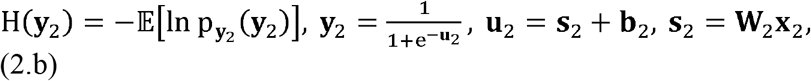

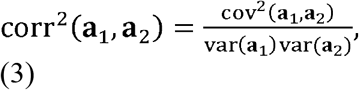

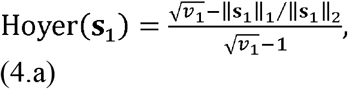

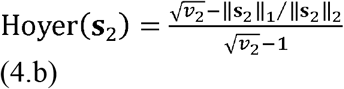

where **X**_1_, **X**_2_ are input data from two modalities with the dimension of subject-by-variable (voxels or SNP loci). Data are modeled as linear mixtures of independent components, i.e., **X**_1_ = **A**_1_**S**_1_,**X**_2_ = **A**_2_**S**_2_. Mixing matrices, **A**_1_ and **A**_2_, have subject-by-component dimensionality and can be estimated as 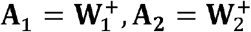, respectively, where ^+^ is the pseudoinverse operation. Independent components, **S**_1_ and **S**_2_, are component-by-variable (voxels or SNP Loci), and can be estimated as **S**_1_ = **W**_1_ **X**_1_ and **S**_2_ = **W**_2_**X**_2_, respectively. Each row of **S**_1_ or **S**_2_ is one independent component (**s**_1_, **s**_2_, respectively), and values in one component reflect the relative involvement of variables (voxels or SNP loci) in the component. Each column of **A**_1_ or **A**_2_ is a loading vector (**a**_1_, **a**_2_, respectively) and their values represent expression levels of the corresponding component across subjects. p_y_1__(**y**_1_) p_**y**_2__ (**y**_2_) are probability density functions of output vectors **y**_1_, **y**_2_, computed as nonlinear transformations of bias-adjusted sources **u**_1_, **u**_2_, respectively. corr(·), cov(·), and var(·) are correlation, covariance, and variance operators, respectively. *v*_1_ and *v*_2_ are the numbers of variables (voxels or SNP loci) in components **s**_1_, **s**_2_, respectively. ||·||_1_ and ||·||_2_ are the *l*_1_ and *l*_2_ norm operators, respectively. λ is a regularizer to balance between independence and correlation maximization. *q*_1_, *q*_2_ are threshold parameters for the Hoyer sparsity index, and 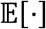 is the expectation operator.

Maximizing the independence of components **s**_1_, **s**_2_ is achieved by maximizing the entropy of output vectors **y**_1_, **y**_2_ using infomax (Bell and Sejnowski, 1995). This is solved by stochastic gradient descent using the relative gradient (Comon and Jutten, 2010) of H(**y**_1_) and H(**y**_2_) with respect to **W**_1_ and **W**_2_, and the traditional gradient of H(**y**_1_) and H(**y**_2_) with respect to **b**_1_ and **b**_2_, respectively (Makeig et al., 1996):

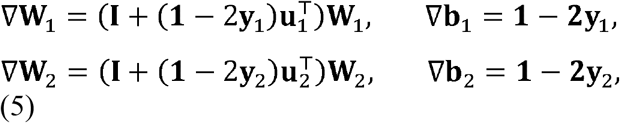

where **I** is an identity matrix, and **1** is a column vector of ones. ⊤ is the matrix/vector transpose operator.

Correlation optimization is solved by gradient descent using the traditional gradient of corr^2^(**a**_1_, **a**_2_) with respect to **A**_1_ and **A**_2_, respectively (Liu et al., 2009):

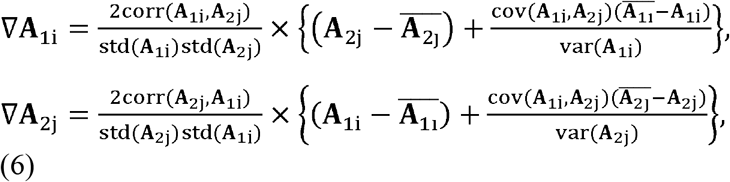

where **A**_1i_ and **A**_2j_ represent the loading coefficients of the i-th component of modality one and the j-th component of modality two, respectively. std(.) denotes the standard deviation, 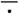 represents an average operation. We set the number of pairs to be optimized as one in this study to avoid overestimating the correlation. So, only the loading coefficient pair with the strongest correlation is optimized for correlation enhancement at each iteration. We monitor the entropy of components (H(**y**_1_) and H(**y**_2_)) during the optimization and reduce λ by a factor of 0.9 whenever the slope of the entropy smaller than – 10^−5^ is detected.

Sparsity constraints are solved by applying the Hoyer projection to the estimated components **s**_1_ = **w**_1_**X**_1_, **s**_2_ = **w**_2_**X**_2_ when their Hoyer indices are less than the preset thresholds *q*_1_, *q*_2_. The Hoyer projection is a nonlinear transformation and is implemented according to the equations above and as described in Hoyer’s paper (Hoyer, 2004). During Hoyer projection, the *l*_2_ norm of the components are unchanged, while the *l*_1_ norm are updated to achieve the desired Hoyer values *q*_1_ and *q*_2_ according to equations 4.a and 4.b. The desired Hoyer index value depends on the data properties of each modality, such as number of variables (voxels or SNP loci), the extent of signal regions, as well as expected minimum signal-to-background ratio (SBR) as defined in (Duan et al., 2019) (the larger the SBR, the easier to separate signal from background). The preset thresholds *q*_1_, *q*_2_ can be estimated from simulated components with an expected (*a priori*) component SBR for each modality, assuming that the background and signal regions of each component form logistic (mean = 0, variance =3) and Laplacian (mean = expected SBR, scale factor = 1) distributions, respectively. Additionally, we conservatively initialize the step sizes for sparsity enhancement (feasibility restoration) to 5 · 10^−3^ (for sMRI) and 5 · 10^−4^ (for SNP) and, in order to prioritize independence maximization, dynamically reduce them by a factor of 0.98 during optimization whenever the slope of the entropy smaller than – 10^−5^ is detected.

Figure 1 shows the flow chart of the proposed spICA, where the dashed boxes and dashed lines indicate extra steps with respect to pICA, and red feedback arrows indicate updates which occur last in each iteration. At the beginning, we initialize the input data **X**_i_, **X**_g_, threshold parameters for the Hoyer sparsity index q_i_ q_g_, learning rates for independence *l*_i_, *l*_g_, incremental step sizes of Hoyer index Δ_hi_, Δ_hg_ for imaging and genetic modalities, separately. Secondly, the unmixing matrices **W**_i_, **W**_g_ are updated based on infomax with the relative gradient of differential entropies **H**(**s**_i_), **H**(**s**_g_) with respect to the unmixing matrices **W**_i_, **W**_g_, respectively. If the Frobenius norm of the difference between the current and previous unmixing matrices is less than a threshold, then the algorithm converges, and the optimization stops for this modality. Otherwise, successively update the component and unmixing matrices using Hoyer projection and correlation optimization, respectively. It is worth noting that pICA optimizes the independence and correlation of components through updates, while sparsity optimization is directly on the components. To take full advantage of sparsity optimization and successfully propagate its effect, we update the data and for the next iteration by reconstructing it from the sparsity-optimized component via product with the pseudoinverse of the current unmixing matrix from infomax. The pseudocode of the proposed spICA is provided in the supplemental materials.

**Fig. 1.**
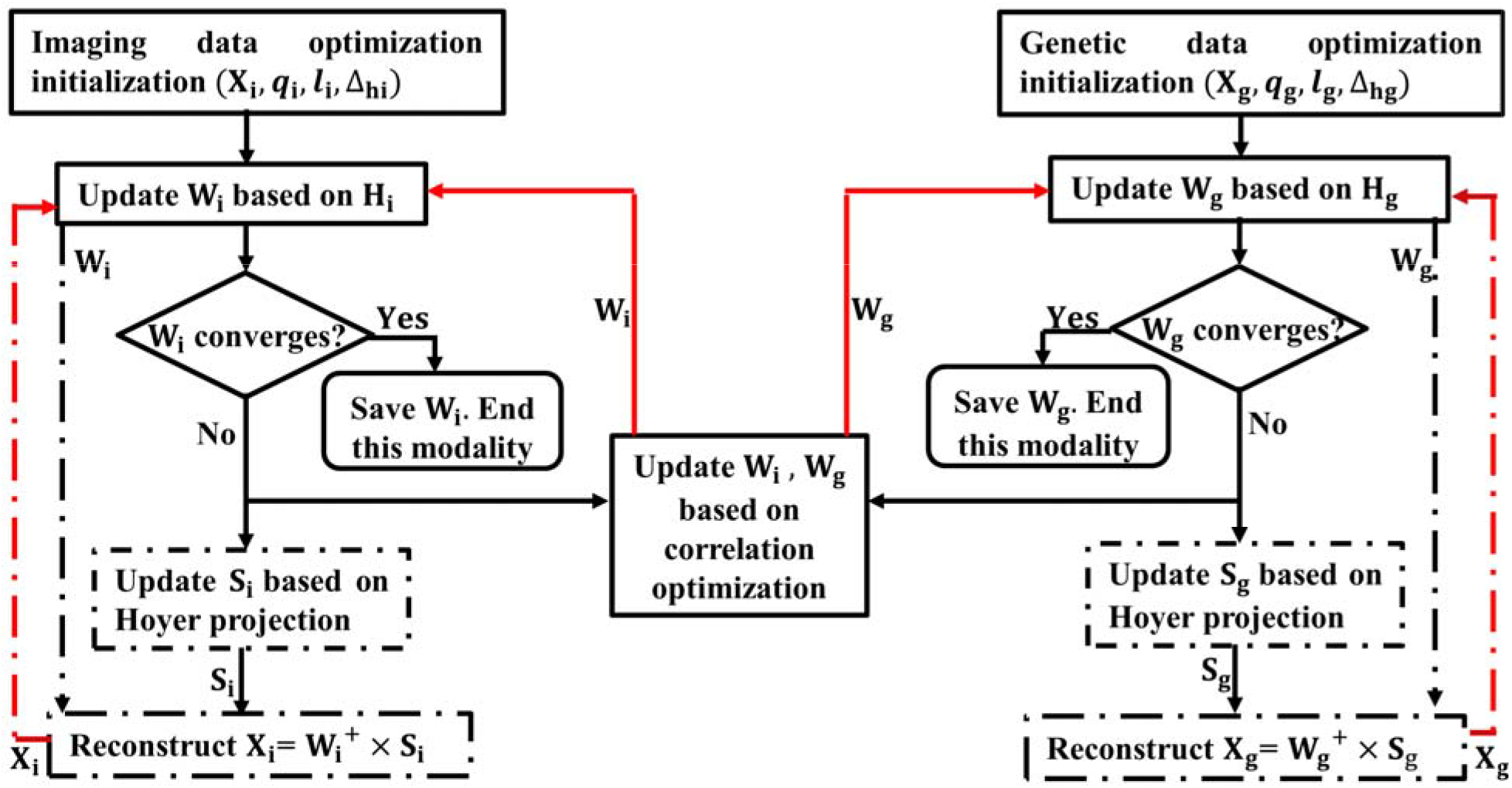
The flow chart of spICA. Note, the dashed boxes and dashed lines indicate extra steps with respect to pICA (red feedback arrows indicate updates which occur last in each iteration).

### 2.2. Simulation design

We evaluated the proposed spICA with simulated structural (s)MRI and SNP data. Its performance was compared with that of pICA under two settings: weak SNP-sMRI correlation (loading correlation of 0.3) and strong correlation (loading correlation of 0.5). Performance measures include the recovered correlation value of the designed sMRI-SNP pair, correlations between recovered sMRI components and ground-truth, correlations between recovered SNP components and designed SNP component patterns, and Hoyer indices of recovered sMRI and SNP components. The regularizer was initialized as one for both spICA and pICA.

SMRI data (dimension: 200 subjects by 31064 voxels) were simulated as the product of a loading matrix (dimension: 200 subjects by 7 components) and a component matrix (dimension: 7 components by 31064 voxels) containing seven non-overlapping brain regions (Figure 2 (a)) generated by the simTB toolbox (Erhardt et al., 2012) (http://trendscenter.org/software/simtb/). All components were designed to have a ground-truth Hoyer index around 0.85 (The Hoyer index value is between 0 and 1. The higher the Hoyer index, the sparser the component). Similarly as in (Duan et al., 2019), the specified Hoyer index was attained by defining the signal regions of components as voxels with intensity larger than 0.25, while the remaining regions were considered as background (the intensities of the background were set to zeros), and the resulting components were used to generate sMRI data. Without loss of generality, the loadings of the *first* sMRI component were designed to be correlated with the loadings of the *first* SNP component (correlation of 0.3 or 0.5 according to the two settings under study). The loadings of the remaining six sMRI components were sampled from a uniform distribution, yielding sample correlations much lower than 0.3 between modalities. White Gaussian noise with a peak signal-to-noise ratio (PSNR) ranging from 7dB to 25dB at a step size of 2dB was superimposed on the component matrix to yield noisy components. The smaller the PSNR, the heavier the noise, thus the more difficult to recover the sources.

**Fig. 2.**
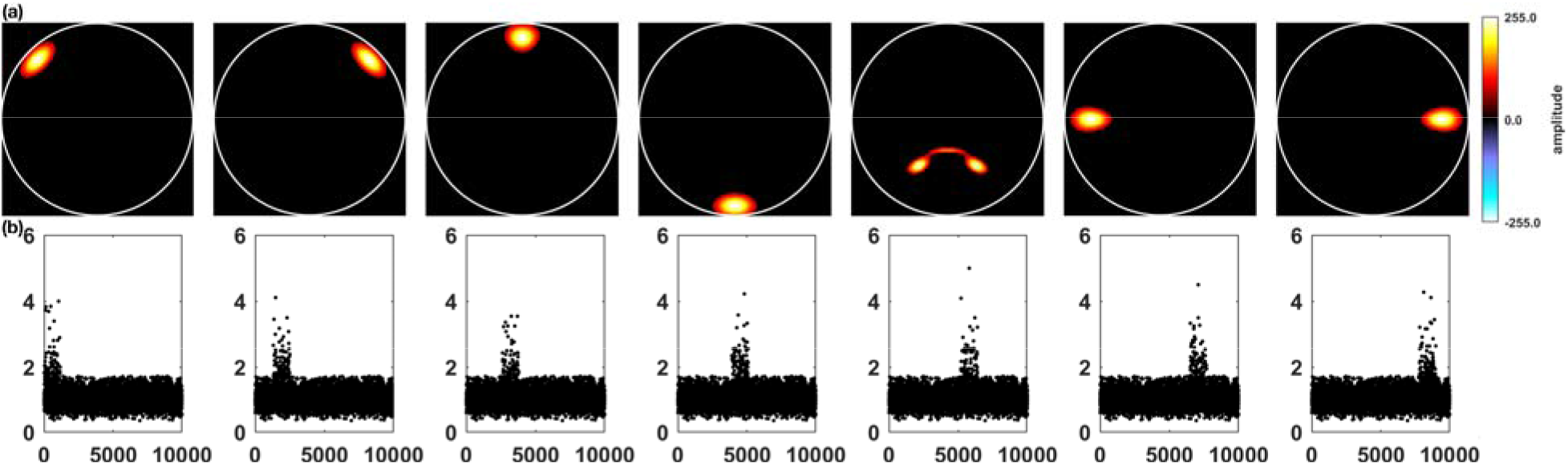
(a) Seven ground-truth sMRI components, (b) The odds ratio of the seven nonoverlapping SNP component patterns.

SNP data (dimension:200 subjects by 10000 SNP loci) were generated using the `simulate` function in PLINK (http://zzz.bwh.harvard.edu/plink/), which were designed to contain 7 nonoverlapping sparse genetic patterns (Figure 2(b)) with corresponding randomly generated case-control status matrix. Each pattern was a realization of one genetic component in the population, where SNPs with higher odds ratio were the SNPs of interest and those with odds ratios around one were background SNPs. All simulated SNPs were unrelated and in linkage equilibrium. For each genetic pattern, 150 loci were set as the SNPs of interest to resemble the sparsity of genetic data. The Hoyer values of such simulated SNP patterns were estimated around 0.4. The case-control status matrix was treated as a proxy of the loading matrix for the SNP modality. Four settings of SNP data were simulated, where the effect size of the patterns varied from 2 to 5 with a step size of one (the corresponding mean odds ratios of the SNPs of interest was between 1.86 to 2.64), and all patterns had the same effect size under each setting (Duan et al., 2019). The effect size is defined as:

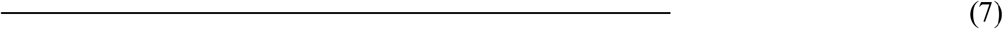

Here, is the representative SNP that best resembled the case-control status and is selected from the set of SNPs of interest (this representative SNP is excluded from the SNPs of interest when estimating the effect size). and denote correlation and standard deviation operations, respectively.

The following scenarios were studied:

**Scenario 1:** Varying noise on the component matrix with fixed sparsity parameters *q*_i_, *q*_g_ for the Hoyer projection procedure in spICA. Different levels of white Gaussian noise (PSNR: 7 dB to 25 dB) were superimposed on the sMRI components, and different levels of effect size (2 to 5) were employed in the SNP patterns. All combinations of PSNR and effect size were evaluated. The sparsity parameters of sMRI and SNP components were fixed to *q*_i_ = 0.85 and *q*_g_ = 0.4 (same as the ground-truth values), respectively.
**Scenario 2:** Fixed noise with varying sparsity settings. White Gaussian noise level of sMRI components was fixed as 21 dB, and the effect size of the SNP pattern was set as 5 (the corresponding mean of odds ratios of the SNPs of interest was around 2.64). Two sets of sparsity settings were tested. (1) The Hoyer constraint threshold of SNP data (*q*_g_) was set as 0.4 (groundtruth value), and the Hoyer constraint threshold of sMRI data (*q*_i_) varied from 0.2 to 0.9. (2) The Hoyer constraint threshold of sMRI data (*q*_g_) was set as 0.85 (ground-truth value), and the Hoyer constraint threshold of SNP data (*q*_g_) varied from 0.2 to 0.9. For each setting under scenarios 1 and 2, spICA was repeated for ten runs. The final value of each performance measure was the average across ten runs, and the corresponding standard error was estimated across ten runs.

### 2.3. Real data analysis

#### 2.3.1. Discovery dataset

The discovery cohort was composed of 341 non-related European Caucasian adults (age: 18-63 years) aggregated from two projects: 198 samples from the NeuroIMAGE project (von Rhein et al., 2015) and 143 participants from the IMpACT-NL sample (Mostert et al., 2015; Onnink et al., 2014), including 127 controls, 167 ADHD patients and 47 unaffected siblings (unaffected siblings and ADHD cases were from different families). Written informed consent was collected from all included samples. All participants were screened to have an IQ larger than 70, and be free from psychosis, addictions in the past six months, current major depression, a diagnosis of autism, epilepsy, neurological disorders, sensorimotor disabilities, as well as any medical or genetic disorders which might be confounded with ADHD (Arias-Vasquez et al., 2019; Onnink et al., 2014; von Rhein et al., 2015). The inclusion criteria for adult ADHD were based on the DSM-IV (NeuroIMAGE project) or DSM-IV-TR (IMpACT-NL). In addition, ADHD assessment in childhood (retrospective for IMpACT-NL, formal research diagnosis for NeuroIMAGE project) was included. Specifically, 18 DSM-IV symptom questions were consistently investigated for all included participants to evaluate their inattention and hyperactivity/impulsivity symptoms. Each symptom outcome ranged from 0 to 9, and the larger the value, the more severe the disorder. In a nutshell, adult ADHD patients had scores larger than 5 in the inattention and/or hyperactivity/impulsivity domain. Unaffected siblings were selected to have a score less than 5 in both inattention and hyperactivity/impulsivity domains. Controls were screened to have a score less than 2 in either symptom domain (Duan et al., 2018). Working memory performance, including maximum digit span forward and backward counts from WAIS Digit Span task (JMR, 2000) were consistently measured for both projects. Significant case vs. control differences have been observed for maximum digit span forward (*p* = 2.62×10^-3^, *t* = 3.02, degree of freedom (DF) = 287) and backward (*p* = 7.27×10^-4^, *t* = 3.96, DF = 287) counts in adults. Table 1 lists the demographics, summaries of cognitive performance, ADHD symptoms, history of stimulants, and scanning sets of the all 341 participants.

**TABLE 1.** Demographics of the adult population

| Variables | Diagnosis group (#) |  |  |
| --- | --- | --- | --- |
|  | Control (127) | Unaffected sibling (47) | ADHD (167) |
| Age | 29.68± 12.35 | 20.91± 2.08 | 25.44± 8.65 |
| Sex(F/M) | 91/36 | 23/24 | 72/95 |
| Estimated IQ | 108.61± 15.36 | 103.60± 12.71 | 103.81± 16.48 |
| Inattention (IA) | 0.52± 1.18 | 1.72± 2.10 | 7.19± 1.74 |
| Hyperactivity<br>/Impulsivity | 0.70±1.00 | 1.51±1.59 | 5.86±2.37 |
| Forward digit span | 9.59± 1.93 | 9.00± 1.74 | 8.89± 1.98 |
| Backward digit span | 7.45± 2.04 | 5.96± 2.10 | 6.42± 2.31 |
| History of stimulants (#) | 1 | 0 | 78 |
| Scan site 1 (#) | 19 | 26 | 52 |
| Scan site 2 (#) | 108 | 21 | 115 |

Note, Scan sites 1 and 2 were used in the NeuroIMAGE project, and scan site 2 was used in the IMpACT-NL project.

#### 2.3.2. Replication dataset

A total of 461 European Caucasian adolescents from 309 families (age: 7-17 years) recruited in the NeuroIMAGE project (von Rhein et al., 2015) were used as the replication dataset, which included 144 controls, 129 unaffected siblings, and 188 individuals fulfilling the diagnosis for ADHD at the time of scanning. Part of the adolescent sample had a sibling in the adult sample. Out of 461 subjects, 452 were over ten years old, and 403 were over twelve years old. Thus, we named this group as adolescents throughout the whole paper. All participants provided their written informed consent. The inclusion and exclusion criteria, evaluations of inattention or hyperactivity/impulsivity symptom, assessments of working memory performance, as well as the grouping criteria for controls were the same as for adults in the aforementioned discovery cohort. The only difference was that adolescent ADHD patients had symptom scores larger than 6 in inattention and/or hyperactivity/impulsivity domain, and unaffected siblings had scores less than 6 in both symptom domains. Significant case vs. control differences for maximum digit span forward (*p* = 1.77×10^-6^, *t* = 4.87, DF = 327) and backward (*p* = 1.90×10^-7^, *t* = 5.32, DF = 327) counts were observed in adolescents. The demographics, summaries of cognitive performance, ADHD symptoms and scanning sets of these 461 adolescents are listed in supplemental Table S1.

#### 2.3.3. sMRI data preprocessing

T1-weighted MRI images for both discovery and replication sets were collected with 1.5T scanners (NeuroIMAGE project utilized Siemens SONATA and Siemens AVANTO, and IMpACT-NL project employed Siemens SONATA) with comparable settings across projects. Exhaustive quality checks have been performed on all T1 images as described previously (Duan et al., 2018; Onnink et al., 2014; von Rhein et al., 2015); only images with good quality were analyzed in this study. Additionally, we computed the coefficient of joint variation (CJV) using MRIQC (Esteban et al., 2017) for all images included. All images had CJV values within three standard deviations from the mean, except for two which had CJV values within four standard deviations away from the mean. Since we consider this amount of variability in CJV reasonable, no images were removed based on the CJV values.

The included T1-weighted images were segmented into six types of tissues (gray matter, white matter, cerebrospinal fluid, bone, soft tissue, and air/background) using SPM 12 (https://www.fil.ion.ucl.ac.uk/spm/software/spm12/) with the default template (for adults) or a customized template (for adolescents) generated by TOM 8 with the matched pair approach (Wilke et al., 2008). Subsequently, gray matter images were normalized into Montreal Neurological Institute (MNI) space using the default spatial normalization function in SPM 12. Normalized gray matter images were then modulated with the Jacobian determinants of the nonlinear transformation and smoothed with a 6×6×6mm^3^ Gaussian kernel. Further quality control, gray matter refinement (see below), and voxel-wise confounding effect (i.e., age, gender, and site, if applicable) correction were performed separately on adults and adolescents. After quality control, only those subjects with a correlation larger than 0.8 with the mean gray matter map were kept (mean gray matter maps for adults and adolescents were generated separately). Gray matter refinement (i.e., masking) selected voxels with mean gray matter volume larger than 0.2 for further analyses, yielding 456921 voxels for adults and 479770 voxels for adolescents (441258 voxels in common for adults and adolescents were used in this study).

Our previous work (Duan et al., 2018; Liu et al., 2020) highlighted that altered gray matter volume (GMV) in three networks, including regions of superior/middle/inferior frontal and cerebellum (ICs 2-4 in Figure S1), consistently related to working memory deficit or inattention in both adults and adolescents. We further confirmed that these regions showed no significant associations with head motion parameters (e.g., CJV, contrast-to-noise ratio, and entropy focus criterion) estimated from MRIQC, indicating a lower likelihood of being driven by head motion.

Identifying the genomic factors underlying these GMV variations was our ultimate goal in the current study. Thus, we reconstructed GMV data of adults and adolescents specifically for these three ROIs (Figure S1, ICs 2-4). Given the whole brain GMV data **X_w_** and the spatial maps S of three ROIs (ICs 2-4 in Figure S1) identified in the previous study (Duan et al., 2018), we computed the projected loading **A_p_** as **A_p_** = **X_w_** × **S**^+^, and then reconstructed GMV data (**X_r_**) as **X_r_** = **A_p_** × **S**, which was then input to spICA. This approach limits the analysis to the portion of the data variance explained by these ROI priors.

#### 2.3.4. Genetic data preprocessing

Both NeuroIMAGE and IMpACT-NL projects used the Illumina Psych Array to genotype DNA extracted from blood. Quality control and imputation were then performed. Samples included in this study fell into a homogenous group (i.e., European ancestry). We further controlled subgroup differences by using five genomic ancestry components in later analyses. Detailed genetic data preprocessing can be found in the supplemental material S1.

A recent large GWAS study on ADHD (Demontis et al., 2019) provides us good prior knowledge to select candidate SNPs for studying genetic factors of brain alterations in ADHD. We selected 2108 SNPs for both discovery and replication datasets by focusing on SNPs showing promising ADHD vs. control difference (*p* < 1×10^-3^) in the GWAS summary statistics (Demontis et al., 2019), and applying a light linkage disequilibrium (LD) pruning (*r*^2^ < 0.9 for p-value informed clumping). Light pruning does not inflate our statistical test, as correlated SNPs end up together in the same component after ICA factorization/decomposition (Chen et al., 2019b).

#### 2.3.5. Analysis of discovery data using spICA

SpICA was performed to decompose 341 independent adults’ reconstructed GMV (**X_r_**, dimension: 341 subjects by 441258 voxels) and SNP data (dimension: 341 subjects by 2108 SNPs) into three independent GMV components and 37 independent SNP components. The GMV component number was set to three due to an expectation that the decomposed GMV components would resemble the three predefined ROIs mentioned at the end of Section 2.3.3 (ICs 2-4 in Figure S1). The regularizer λ was initialized as one. No sparsity constraint was imposed on GMV data since signals can be easily separated from backgrounds for these three ROI priors (we set *q*_i_ = 0 so that spICA would not attempt to optimize the sparsity for GMV data). The Hoyer constraint threshold of SNP data was set as 0.4 (i.e., *q*_g_ = 0.4). The SNP component number was estimated based on Chen’s consistency measure (Chen et al., 2012). SpICA was performed ten times, and ICASSO (Himberg and Hyvarinen, 2003) was employed to select the most stable run to calculate GMV-SNP associations. Bonferroni correction was applied at *p* < 0.05 for comparison of 3*37 GMV-SNP associations.

Moreover, spICA was performed on 100 subsampled sets to test the stability of the identified GMV-SNP pairs, where stratified sampling was employed to randomly select 90% subsamples from each of the ADHD, control, and unaffected sibling groups. To test the likelihood of overfitting for the identified GMV-SNP associations, 1000 random permutations of the subjects were performed separately in the GMV and SNP data. SpICA was then applied to the permuted data, and a null distribution was obtained based on the resulting pairwise associations (i.e., the subject expression level correlations). Then a p-value was computed as the percentage of pairs yielding significant GMV-SNP associations (Bonferroni correction was applied at *p* < 0.05 for 1000*37*3 pairs), which reflected the probability of overfitting of the spICA model.

We further checked the stability of the identified SNP component under two scenarios: (1) the number of SNP component varied from 5 to 40 (SNP data were fixed with *p* < 1×10^-3^, *r*^2^ < 0.9 preselection); (2) SNP preselection *p*-value varied from 10^-4^ to 10^-2^ (*r*^2^ < 0.9) and the number of SNP component was estimated for each resulting SNP data according to Chen’s consistency measure (Chen et al., 2012), see supplemental material S2 for detail. The stability of the identified GMV-SNP associations was examined by performing spICA on a heavily pruned SNP data (*r*^2^ = 0.2, *p* < 1×10^-3^ preselection).

#### 2.3.6. Validation of GMV-SNP component associations in the replication dataset

The replication dataset contained GMV and SNP data of 461 adolescents. Given that the discovery and replication datasets included participants of different age groups (the discovery dataset included adults and the replication dataset included adolescents), the strongest linked GMV-SNP patterns may be different for these two population (no matter what fusion methods (e.g., sCCA, pICA, or spICA) were applied). That is because all these association-driven methods were designed to optimize the correlation between modalities. Thus, the pair identified in the discovery dataset might even be absent in the replication dataset. This could cause discovery components to be undetectable in the replication set using association-driven fusion approaches. Thus, to investigate the identified discovery GMV-SNP pair in the replication dataset, we used a projection method with the assumption that the replication dataset shared the same components. The following projection method is utilized: let **S_di_** and **S_dg_** denote the source/component matrices of adults’ (discovery) GMV and SNP data, respectively. Let **X_ri_** and **X_rg_** represent adolescents’ (replication) GMV and SNP data, respectively. Then the corresponding loading matrix of the adolescents’ GMV data can be estimated as 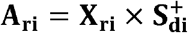. Recall that spICA utilized nonlinear sparsity regularization and reconstructed SNP data with cleaner sources at each iteration (i.e. denoising the SNP components of the discovery dataset). Thus, we reconstructed the loading matrix of SNP data using Tikhonov-regularized least squares (Strang, 2007) to account for the denoising effect in the replication SNP data. The loading matrix of SNP data was reconstructed as 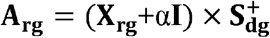, where **I** is an identity matrix, α is a parameter calculated as in (Duan et al., 2019) to balance between retaining the signal and reducing the noise. Associations between GMV and SNP loadings for the identified pairs were assessed in the replication dataset using linear mixed effect models with family as a random effect, as explained in the following section. The identified GMV-SNP pair was considered as replicated if the corresponding corrected p value was less than 0.05 in the replication dataset.

To investigate whether participants from different age groups presented similar GMV-SNP associations within the 317 adolescents from ADHD families (i.e., 188 cases and 129 unaffected siblings) in the replication dataset, we evaluated GMV-SNP associations in five subgroups partitioned by age distribution (as listed in Table 2), separately.

**TABLE 2.** Subgroups partitioned by age of 317 adolescents from ADHD families in the replication sample.

|  | Age Range, # | ADHD (#) | Unaffected Siblings (#) | Female (#) | Male (#) |
| --- | --- | --- | --- | --- | --- |
| Partition 1 | 7-15, 152 | 96 | 56 | 81 | 71 |
|  | 15-17, 165 | 92 | 73 | 62 | 103 |
| Partition 2 | 7-14, 108 | 68 | 40 | 65 | 43 |
|  | 14-16, 91 | 46 | 45 | 35 | 56 |
|  | 16-17, 118 | 74 | 44 | 43 | 75 |

Note, # denotes the number of subjects.

#### 2.3.7. Statistical association analyses

For the discovery data (adults’ data), the associations among GMV components, SNP components, working memory, and symptom scores were examined using the following linear regression models:

1. a GMV component loading = a SNP component loading + five genomic ancestry components.
2. working memory or symptom variable = age + gender + a GMV component loading.
3. working memory or symptom variable = age + gender + a SNP component loading + five genomic ancestry components. In model 1, each GMV/SNP component was tested separately. Age and gender effects were not considered for GMV components in model 1 because they have been regressed out voxel-wisely in the preprocessing step (the same for models 5 and 6 below). Working memory performance was measured by maximum forward and backward digit span count. The symptom variable included inattention and hyperactivity/impulsivity symptoms. The case vs. control difference of each GMV component was evaluated using a two-sample t-test, and case vs. control difference of each SNP component was assessed using the following model:
4. a SNP component loading = diagnosis (case/control) + five genomic ancestry components. For the replication data (adolescents’ data), family structure and medication history (binary values) were considered in the association analyses, as medication did affect the three brain ROIs studied here in adolescents included in this study (Liu et al., 2020). Thus, the association between a GMV component and a SNP component and the case vs. control difference for a GMV/SNP component were tested using the following linear mixed effect models:
5. a GMV component loading = a SNP component loading + five genomic ancestry components + family ID + medication status.
6. a GMV component loading = diagnosis (case/control) + medication status + family ID.
7. a SNP component loading = diagnosis (case/control) + family ID + five genomic ancestry components.

In models 5-7, the family structure was modeled as a random effect, and other predictors, i.e., a SNP component loading, diagnosis, medication, and five genomic ancestry components, were treated as fixed effects.

Confounding effects, including major depression and IQ (medication and anxiety were also treated as potential confounding factors for adults in models 1-3), were examined for both discovery and replication data by adding them as a covariate, one at a time, to the models above.

#### 2.3.8. Interpretation of identified GMV and SNP components

Considering that signs of components from ICA are arbitrary, we assigned a sign for each identified GMV/SNP component so that the peak value was positive, and then adjusted the sign of the corresponding loadings accordingly. Each of the identified GMV and SNP components were further normalized to have a zero mean and unit standard deviation. GMV component regions were selected with |*z*| > 2.5 and mapped into the Talairach atlas (Lancaster et al., 2000). SNPs with absolute weights larger than 2 were selected for further interpretation of gene annotation, enrichment analysis, potential expression quantitative trait loci (eQTL) and methylation quantitative trait loci (mQTL) effects. Moreover, we performed univariate association analysis between individual SNP loci and loadings of the identified GMV components, controlling for five genomic ancestry components, and reported in the Manhattan plot of the univariate association p-values, which was compared to the identified SNP component via visual inspection.

## 3. Results

### 3.1. Simulation

The effectiveness and robustness of spICA were examined under two scenarios: 1) varying noise level in sMRI components and varying effect size in SNP data, and 2) varying the sparsity parameters *q*_i_, *q*_g_ for extraction of sMRI and SNP components, respectively.

Figure 3 and 4 present results under scenario 1 with the ground-truth correlation of 0.3 (similar results were obtained for ground-truth correlation of 0.5 in Figure S3 and S4). Figure 3 displays the recovered sMRI-SNP correlations. Recall that the sparsity parameters of spICA were fixed to match the ground-truth values: *q*_1_ = 0.85 (sMRI) and *q*_g_ = 0.4 (SNP). As expected, performances from both spICA and pICA indicated that the larger the effect size of SNP data and the larger the PSNR value, the more accurate the detected sMRI-SNP association. The sMRI-SNP association accuracy is fairly stable with respect to PSNR, dropping with PSNR < 9. Under most conditions (i.e., the effect size of SNP data from 3 to 5), the sMRI-SNP correlations identified by spICA were much closer to the ground-truth value of 0.3, as compared to those from pICA. When the effect size of SNP data was 2 (corresponding odds ratio of 1.86), both spICA and pICA failed to detect the designed association.

**Fig. 3.**
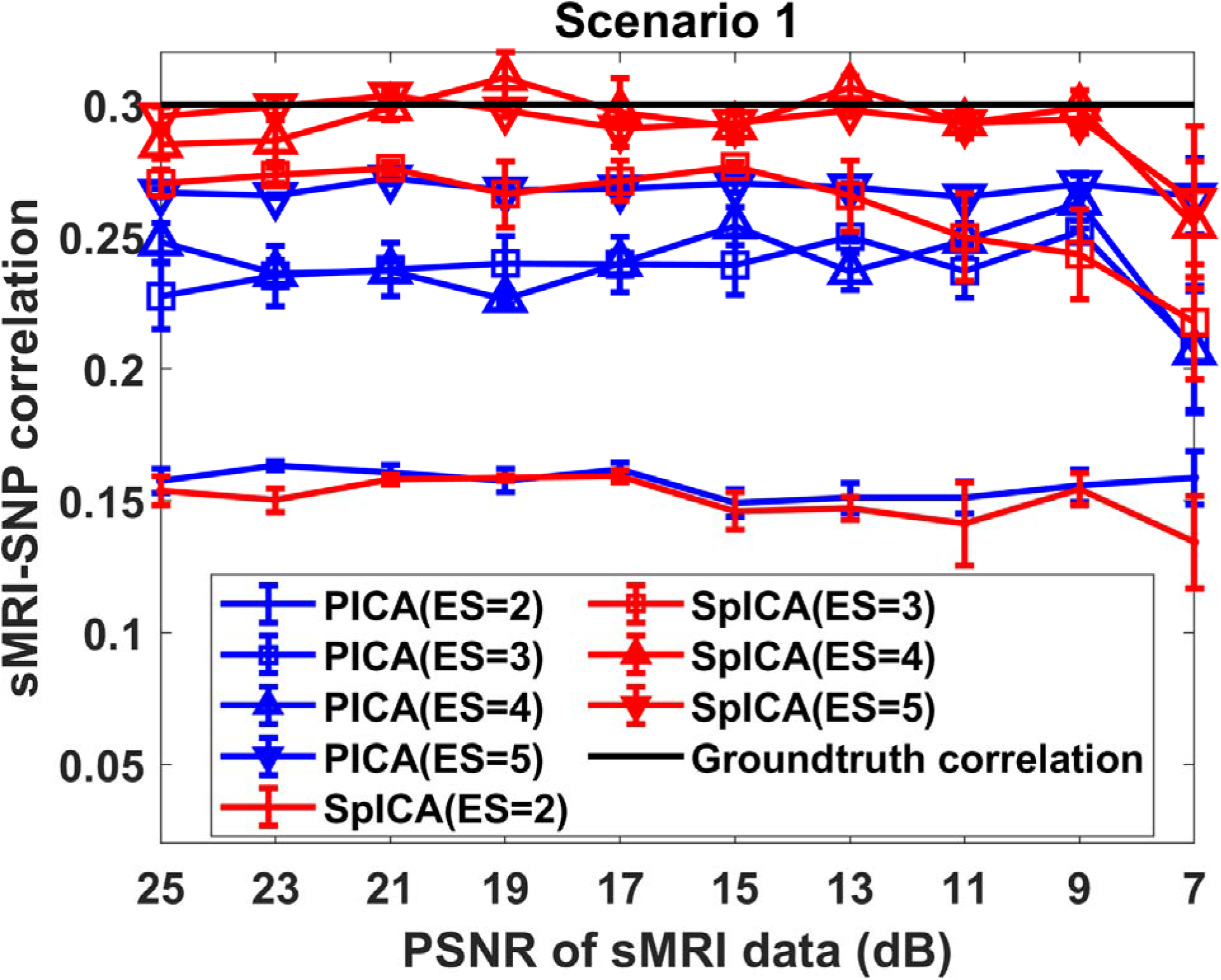
Scenario 1: The sMRI-SNP correlations detected by spICA (red) and pICA (blue) while varying noise levels of sMRI components and effect size of SNP data. The ground-truth of the correlation was 0.3 (black). The markers dot, square, upward triangle, and downward triangle represent results for SNP data with an effect size (ES) of 2, 3, 4, and 5, respectively. Note, color representations are consistent for all simulation results (Fig. 3–5 and Fig. S3-S5), and shape representations are consistent for the sMRI-SNP correlation of 0.5 (Fig. S3).

**Fig. 4.**
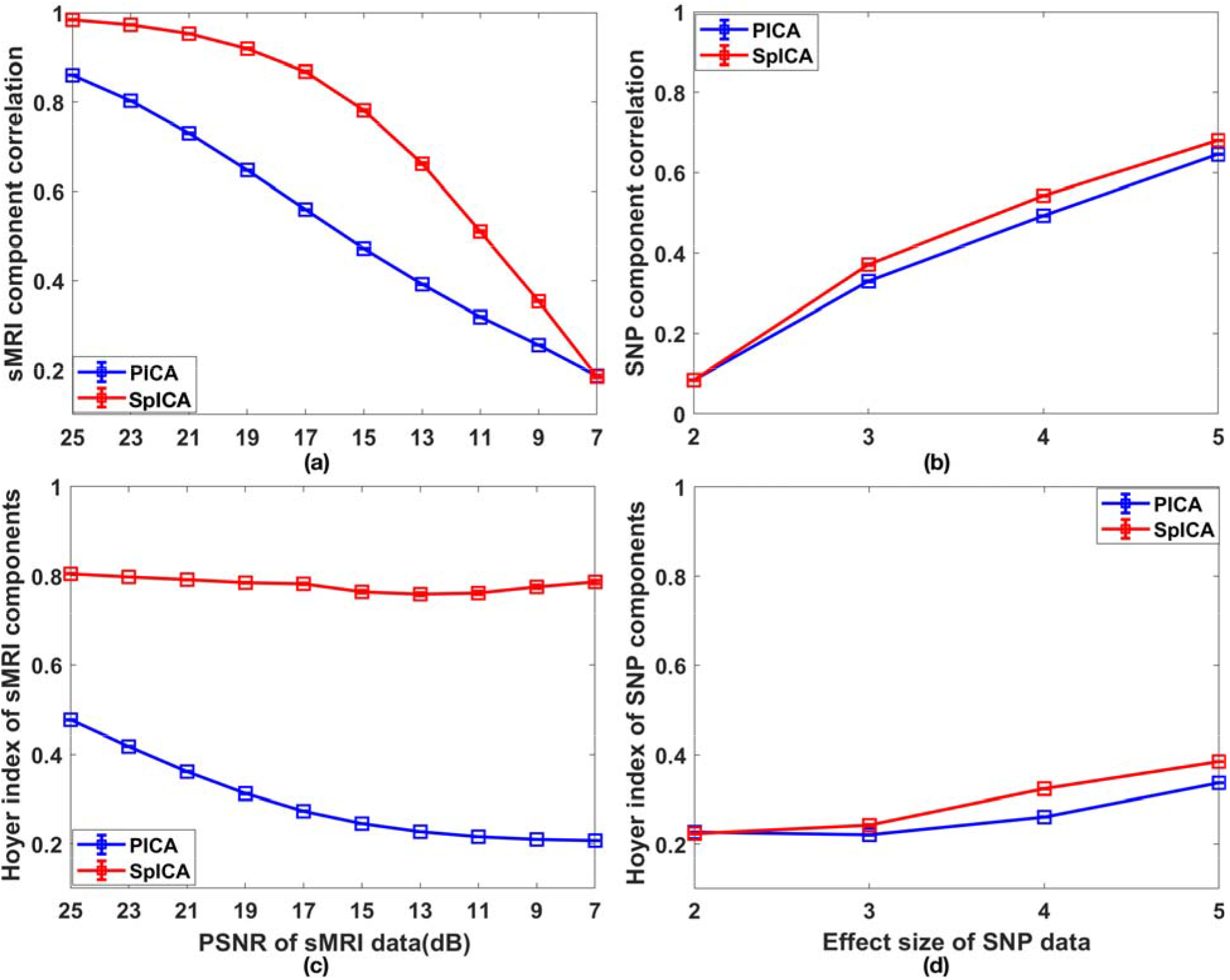
Performance of spICA and pICA while varying PSNR values of white Gaussian noise superimposed on sMRI components and effect size of SNP data: (a) accuracy and (c) sparsity of recovered sMRI components; (b) accuracy and (d) sparsity of recovered SNP components (the designed sMRI-SNP association was fixed at 0.3).

The accuracy of sMRI and SNP components (correlations with ground-truth components), and the sparsity of sMRI and SNP components (Hoyer indices) are displayed in Figure 4. The detection accuracy and Hoyer indices of sMRI components plotted in Figure 4 (a) and (c) are the results across different SNP effect size settings. Similarly, Figure 4 (b) and (d) plot the detection accuracy and Hoyer indices of SNP components across different PSNR settings. Varying the effect size of SNP data has a negligible influence on sMRI component recovery. Similarly, varying the PSNR of sMRI components has a trivial influence on SNP component recovery (negligible error bars in the plots). Figure 4 (a) and (c) demonstrated that as PSNR decreased, the accuracy of sMRI components decreased for both spICA and pICA, while the sparsity of sMRI components decreased for pICA but not so much for spICA. Nonetheless, spICA always yielded higher recovery accuracy and sparsity compared to pICA when PSNR was larger than 7. Figure 4 (b) and (d) indicate that as the effect size of SNP data increased, accuracy and sparsity of SNP components increased for both spICA and pICA. In addition, spICA outperformed pICA in terms of both accuracy and sparsity when the effect size of SNP data was larger than 2.

For scenario 2, with a designed sMRI-SNP correlation of 0.3 and various sparsity constraint threshold settings (*q*_i_, *q*_g_), the recovered sMRI-SNP association, sMRI and SNP components accuracy, and sparsity indices from both spICA and pICA were plotted in Figure 5 (similar results were obtained for sMRI-SNP association of 0.5, see Figure S5). When *q*_i_, *q*_g_ were less than 0.3 (largely inactive constraints) for sMRI and SNP data, respectively, spICA and pICA presented a similar performance for sMRI-SNP association detection and sMRI and SNP components detection. As *q*_i_, *q*_g_ increased and approached the Hoyer index values of groundtruth components, the identified sMRI-SNP association gradually approached the designed association of 0.3, and accuracies of sMRI and SNP components improved as well. When *q*_i_ for sMRI data exceeded the ground-truth Hoyer value (i.e., 0.85), accuracies of recovered sMRI components dropped but remained higher than those from pICA. When *q*_g_ for SNP data exceeded 0.5, accuracies and sparsity measures of the recovered SNP components remained largely unchanged. This was likely due to smaller feasibility restoration step sizes for the Hoyer projection of SNP data at each iteration, resulting from our strategy to prioritize independence via monitoring entropy (see Section 2.1) and causing the restoration step to become small. This has the effect of avoiding both abrupt changes to the SNP data and deterioration of the components’ independence. It also means that the sparsity constraint is enforced only insofar as the data supports statistical independence. In sMRI components, note that higher noise levels dramatically decrease sparsity. Thus, a relatively large increment in Hoyer sparsity (via larger feasibility restoration step sizes) is needed to enhance the sparsity of sMRI components effectively. This increment in sparsity is less likely to deteriorate the component’s independence because here noise occurs at the component level.

**Fig. 5.**
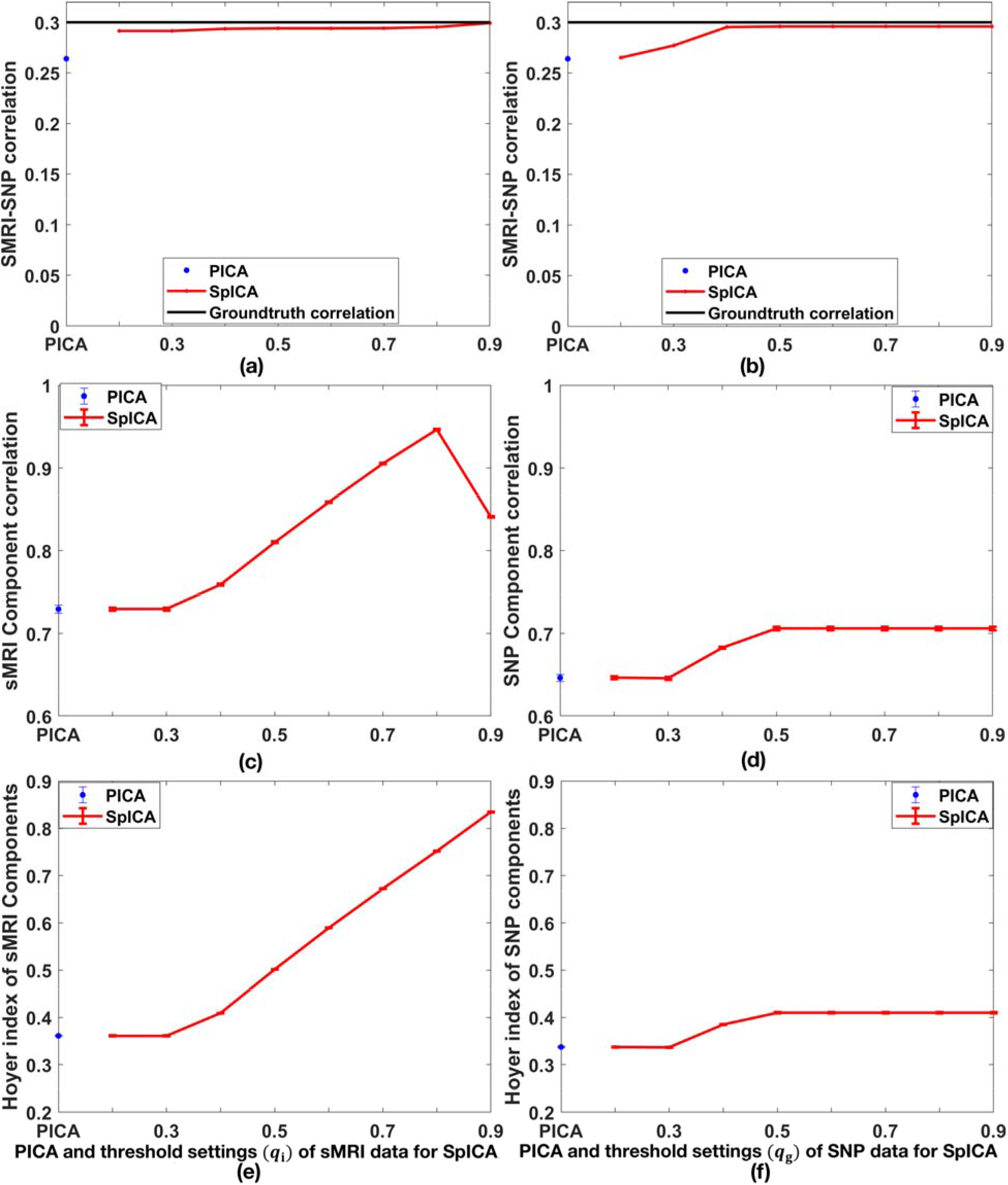
Scenario 2: sMRI-SNP association detections from pICA and spICA with PSNR of 21dB and effect size of 5, varying Hoyer index thresholds from 0.2 to 0.9 for (a) sMRI data (varying with fixed, the ground-truth value for SNP data), (b) SNP data (varying with fixed, the ground-truth value for sMRI data); corresponding spICA and pICA performances including (c) accuracy and (e) sparsity of recovered sMRI components while varying from 0.2 to 0.9; corresponding spICA and pICA performances including (d) accuracy and (f) sparsity of recovered SNP components while varying *q*_g_ from 0.2 to 0.9 (the designed sMRI-SNP association was fixed at 0.3).

### 3.2. Real data analysis

#### 3.2.1. Discovery results

For the discovery data, spICA identified that two GMV components (IC 1 and IC 2 in Figure 6 (a) and (b)) showed significant and positive associations with one SNP component in Figure 6 (c) (IC 1-SNP_*p* = 6.41×10^-13^, 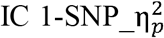 (partial eta squared) = 0.35; IC 2-SNP_*p* = 2.46×10^-4^, 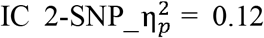) after controlling for population structure using five genomic ancestry components and applying Bonferroni correction (0.05/(3*37)). Among 100 stratified subsampling tests, 89 subsamples stably identified GMV IC 1-SNP and GMV IC 2-SNP pairs with similar correlation strengths as in the full sample. Among all pairwise correlations (i.e., 1000*3*37 pairs in total) computed from 1000 permutation test, 76 passed Bonferroni correction, yielding to a tail probability of *p* = 6.85×10^-4^ for the null distribution. Based on the subsampling and permutation results, we can infer that the identified GMV IC 1-SNP and GMV IC 2-SNP associations were stable, and the corresponding correlation coefficients were likely not overfitted. Positive associations between GMV ICs 1-2 and the identified SNP component indicated that more counts of minor/reference alleles in SNPs with positive weights (hot colored loci in Figure 6 (c)) were related to higher GMV in brain regions with positive weights (hot colored regions in Figure 6(a) and (b)), and vice-versa. Figure 6(d) and (e) show the associations between loadings of the SNP component and loadings of GMV IC 1 and IC 2, respectively, where scatter plots of controls, cases, and unaffected siblings are in red, green and blue colors, respectively (the same for Figure 6(f)). We observed that the GMV IC 1-SNP component association was much stronger in unaffected siblings and ADHD patients compared to controls 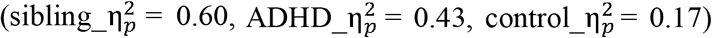. Further, we treated subjects from ADHD families (i.e., cases and unaffected siblings) as one group and participants from control families (i.e., controls) as another group. We then observed that this categorical (binary) family label significantly interacted with SNP loading for GMV IC 1 prediction (*p* = 8.50×10^-3^) after extending model 1 (Section 2.3.6) to GMV IC 1 loading = SNP loading + ancestry + family label + family label*SNP loading. Medication, IQ, and comorbidity did not confound the identified GMV-SNP associations.

**Fig. 6.**
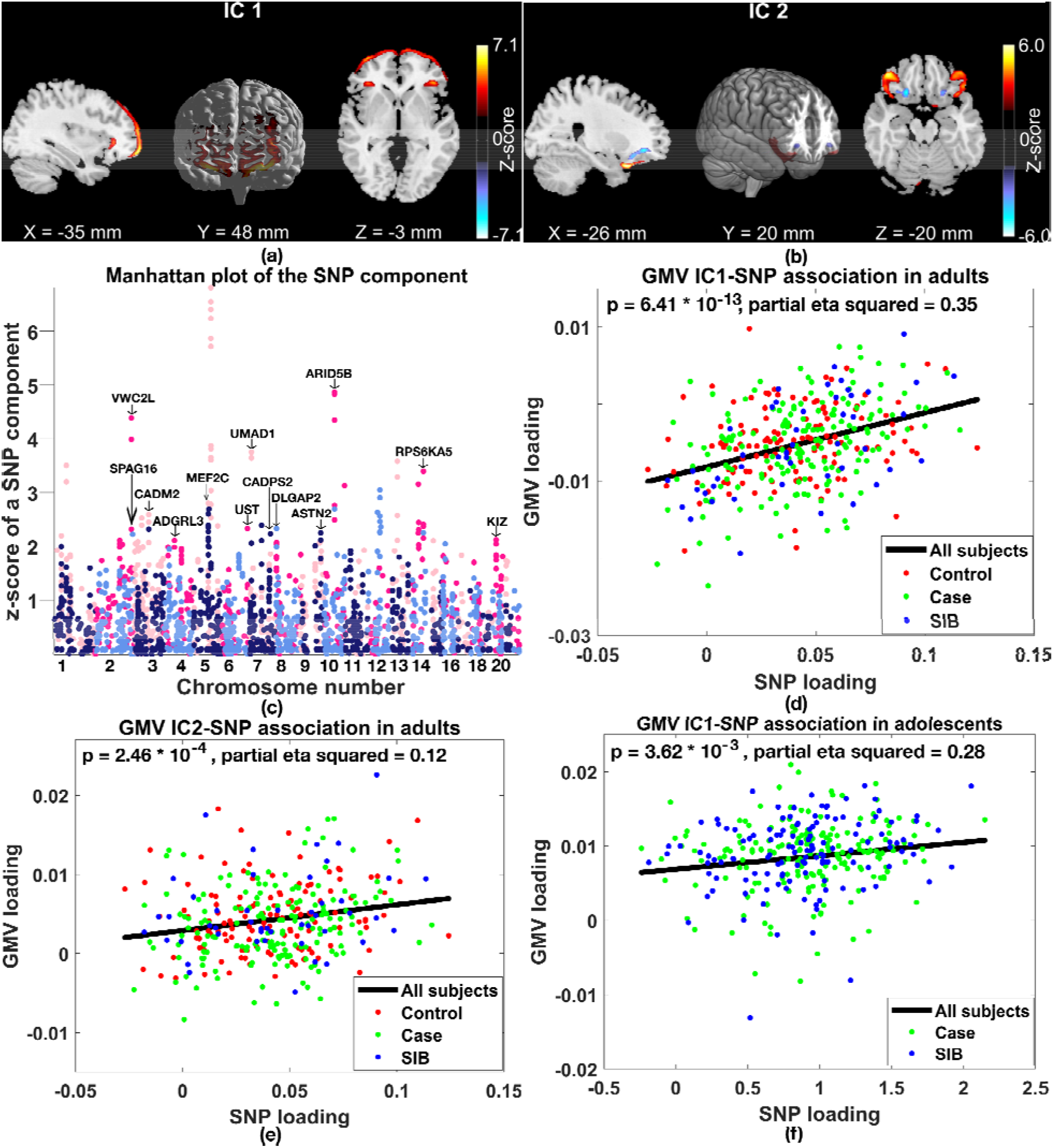
SpICA results in discovery and replication sets. (a) GMV IC 1, (b) GMV IC 2, (c) the identified SNP component, the association between GMV IC 1 and SNP component (d) in adults (the discovery set) and (f) adolescents (the replication set), (e) the association between GMV IC 2 and SNP component in adults. In subplots (a)-(c), hot colors represent positive weights, and cold colors denote negative weights. In subplots (d)-(f), black, red, green, and blue colors represent all subjects, controls, cases, and unaffected siblings (SIB).

We confirmed that the identified SNP component was robust to the total number of components decomposed for SNP data (SNP component number range: [24, 40]) and SNP preselection p values (p: [0.0005, 0.005])--additional details in supplemental material. Applying spICA to the heavily pruned SNP data (*r*^2^ = 0.2, *p* < 1×10^-3^), we identified a similar pair as reported in the main finding (the GMV IC 1-SNP pair), indicating that the discovered GMV IC 1-SNP association was unlikely biased by LD structure. See supplemental material S2 for details.

#### 3.2.2. Replication results

The association between GMV IC 1 and the SNP component was nominally significant in 461 adolescents after controlling for medication and five genomic ancestry components (uncorrected *p* = 4.60×10^-2^, 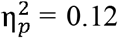), where loadings of GMV and SNP components were computed using the projection method as described in section 2.3.6. The association between GMV IC 2 and the SNP component was not replicated in 461 adolescents. Similar as in the discovery set, we observed a much stronger association for GMV IC 1-SNP component in unaffected siblings and ADHD patients compared to controls 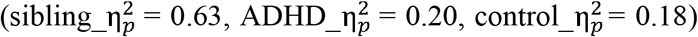. Focusing on adolescents from ADHD families (i.e., 317 adolescents, including 129 unaffected siblings and 188 cases), the association between GMV IC 1 and SNP component became significant (Figure 6 (f), corrected *p* = 7.24×10^-3^, 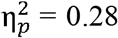, corrected for two GMV-SNP pairs identified in the discovery samples); the association between GMV IC 2 and SNP component remained not significant. IQ and major depression did not confound the association between GMV IC 1 and the SNP component in adolescents from ADHD families. Moreover, 165 older adolescents (age: 15-17 years composed of 92 cases and 73 unaffected siblings) demonstrated significant positive GMV IC 1-SNP component association (uncorrected *p* = 2.19×10^-2^, 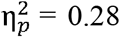), while it was not significant in younger ones (age: 7-15 years, *p* = 1.75×10^-1^). The GMV IC 2-SNP component association was not significant in both age groups (i.e., age range: 7-15 years and age range: 15-17 years). Similarly, GMV IC 1-SNP components association was nominal significant in the 118 oldest adolescents (age: 16-17 years, uncorrected *p* = 3.71×10^-2^, 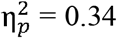, beta > 0) but not in younger ones (age: 7-14 years and age: 14-16 years). Moreover, GMV IC 2-SNP component association was nominal significant in the 118 oldest adolescents (age: 16-17 years, uncorrected p = 4.65×10^-2^, 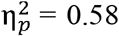, beta > 0) but not in younger ones (age: 7-14 years and age: 14-16 years).

#### 3.2.3. Brain region identification

The identified GMV IC 1 (z-scored) is illustrated in Figure 6(a), highlighting superior and middle frontal gyri (|z| > 2.5). The peak of GMV IC 1 had a value of z = 7.1 and was located at x = −22, y = 60, z = −16 (MNI coordinates). For loadings of GMV IC 1, we did not observe significant GMV differences between individuals with ADHD and control in adults, while we observed significant GMV reduction for those with ADHD in adolescents after controlling for the medication effect (*p* = 8.72×10^-3^, *t* = 2.64, DF = 329). Loadings of GMV IC 1 significantly and positively associated with backward (*p* = 2.09×10^-2^, 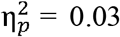, beta > 0) and forward (*p* =2.41×10^-2^, 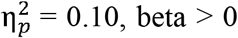, beta > 0) digit span performance in adults and adolescents, respectively. Brain regions of GMV IC 2 are described in supplemental material S3.

#### 3.2.4. Highlighted SNPs identification, their annotations, and regulatory effects

Figure 6(c) shows the absolute values of the z-scored SNP component associated with GMV ICs 1-2, where positive weights are displayed with hot colors (i.e., pink and deep pink), and negative weights were plotted with cold colors (i.e., midnight blue and cornflower blue). Hot/cold colors with two levels of tint distinguish two sequential chromosomes. No significant case vs. control difference was observed for loadings of the identified SNP component in both discovery and replication sets. Loadings of the identified SNP component were not significantly associated with forward/backward digit span performances or symptom scores.

The identified SNP component highlighted 93 top SNPs (|z| > 2). Using human hg 19 build, 93 top SNPs were mapped to genes encoding long non-coding RNAs and 24 protein-coding genes, which were not significantly enriched in any pathways from Gene Ontology analysis. Performing Ingenuity Pathway Analysis (IPA) on the 24 protein-coding genes, five genes, including MEF2C, DLGAP2, CADPS2, CADM2, and ADGRL3, were identified being involved in cell-to-cell signaling and interaction (*p*-value range: [2.24×10^-4^, 4.18×10^-2^], molecules number: 5). Six genes, i.e., MEF2C, DLGAP2, CADPS2, CADM2, ADGRL3, and UST, played a role in the nervous system development function (*p*-value range: [2.24×10^-4^, 4.18×10^-2^], molecules number: 6). In addition, MEF2C, DLGAP2, and CADPS2 exerted effects in excitatory postsynaptic potential (*p* = 2.24×10^-4^, molecules number: 3).

The top SNPs do not significantly regulate the expression of genes in Brodmann BA 9 region based on the Genotype-Tissue Expression (GTEx) database (Ardlie et al., 2015). While a top SNP, rs13166522, presented a nominal regulation effect on the expression of gene NUDT12 (*p* = 1.32×10^-2^) in the Brodmann BA 9 region (Ardlie et al., 2015). Focusing on Ramasamy’s (Ramasamy et al., 2014) summary on eQTL in the frontal cortex with a nominal *p* < 10^-3^, 10 top SNPs regulated expression of 10 genes including NUDT12, RHOJ, as listed in Table S3. Out of 93 top SNPs, ten significantly regulated methylation levels of 10 CPG sites (see Table S4 for detail) using mQTL in the human frontal cortex provided in (Jaffe et al., 2016).

#### 3.2.5. Genetic association analysis of the GMV loading

Figure 7 displays the Manhattan plot of −log_10_(p) of the association between each SNP and loadings of GMV IC 1 in 341 adults, where interchange pink and dark pink colors were employed to distinguish two sequential chromosomes. After FDR correction, none of the 2108 SNPs showed a significant association with GMV IC 1 (the same for the SNP dataset with heavy pruning of *r*^2^ = 0.2). The p-value map of the association highlighted SNP loci in chromosomes 5, 1, 2, 3, 7, and 14, which were in line with the highlighted regions in the identified SNP component (Figure 6(c)) based on visual inspection.

**Fig. 7.**
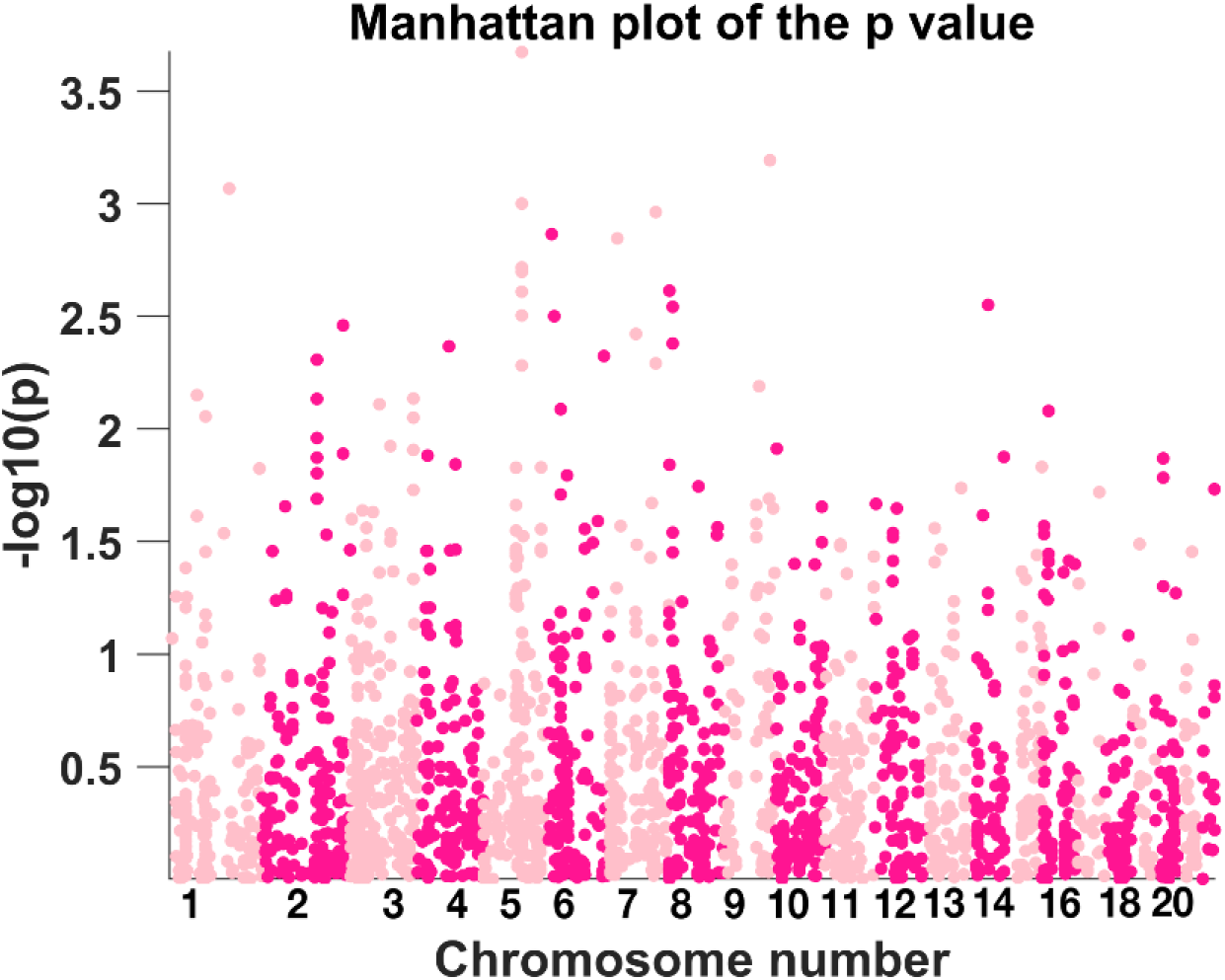
Manhattan plot of −log_10_(p) of the association between individual SNP and loadings of GMV IC 1 in adults.

## 4. Discussion

In this study, we proposed a multivariate fusion method, spICA, for imaging-genetic studies. While building upon pICA for simultaneous independence maximization (via infomax ICA) and correlation optimization, spICA seeks further improvement in performance by leveraging the sparsity nature of sources. The sparsity constraint is implemented through a nonlinear Hoyer projection, which is applied directly to the estimated sources to suppress noise (background) and enhance the signal. Additionally, the sparsity enhancements are continually propagated by passing the reconstructed, cleaner data to the next iteration, yielding a net-nonlinear transformation overall.

Simulation results demonstrate that spICA outperforms pICA with improved accuracy for sMRI-SNP association detection and component spatial map recovery, as well as with enhanced sparsity for sMRI and SNP components under both strong (*r* = 0.5) and weak association (*r* = 0.3) settings. As illustrated in the Methods section, preset Hoyer constraint thresholds can be estimated based on the components simulated with expected SBR and the amount of signal to retain. Simulations suggest that overestimating the ground-truth Hoyer values (ergo, setting a higher constraint threshold) yields decreased accuracy for spatial maps recovery in sMRI data. That is not the case for SNP data, suggesting that a moderate overestimation of the preset Hoyer constraint threshold for SNP data is acceptable.

Applying spICA to GMV and SNP data of 341 unrelated European Caucasian adults, we identify one SNP component (Figure 6(c)) as significantly and positively associated with two GMV components (GMV ICs 1-2 in Figure 6(a-b)). Moreover, the association between SNP and GMV IC 1 is replicated in 317 adolescents from ADHD families. This association interacts with the family categories (i.e., case and control families) in both adults (*p* = 8.50×10^-3^) and adolescents (*p* = 3.68×10^-2^), i.e., it is significant in subjects from case families but not in those from control families. Additionally, GMV IC 1-SNP association is stronger in adults 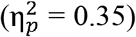 compared to adolescents 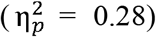. Within 317 adolescents from ADHD families, GMV IC 1-SNP association is stronger in older participants (age: 15-17 years or 16-17 years) compared to younger ones (age: 7-15 years or 7-14 years, or 14-16 years); GMV IC 2-SNP association is nominal significant in older subjects (age:16-17 years) but not in younger ones (age: 7-14 years or 14-16 years), indicating that this association is likely adult-specific. These findings suggest that participants from ADHD families carrying higher loading value of this SNP component presented higher gray matter volume in the superior and middle gyri. This relation became stronger with age. Interestingly, for GMV in superior and middle gyri (GMV IC 1), ADHD patients showed no significant difference compared to controls in adults but demonstrated significant GMV reduction in adolescents. Collectively, these findings lend support to the theory that ADHD patients present delayed development in the frontal cortex (Krain and Castellanos, 2006; Rubia, 2007; Shaw et al., 2007; Shaw et al., 2012), suggesting that top SNPs of the identified SNP component likely compensate GMV reduction in the frontal cortex in ADHD patients, and this compensation effect may have a late onset.

Recently, Hoogman, et al. (Hoogman et al., 2019) conducted a meta-analysis on the large-scale ENIGMA ADHD samples to characterize subcortical volumes and cortical thickness/surface area alterations across the lifespan between individuals with ADHD and controls using a split-half validation method. They highlighted that widespread cortical surfaces, including the total surface and superior frontal surface, presented significantly reduced area in children with ADHD, and the reduction was attenuated in adolescents and adults with ADHD. Later, Zhao and colleges (Zhao et al., 2020) characterized GMV reduction in left superior frontal and right middle frontal gyri in ADHD adolescents using sMRI data from ADHD 200 project. Altogether, our results are in line with these findings showing that GMV reduction in superior and middle frontal gyri occurs in ADHD children/adolescents but diminishes in ADHD adults, suggesting a delayed maturation in frontal cortex (Krain and Castellanos, 2006; Rubia, 2007; Shaw et al., 2007; Shaw et al., 2012). Better working memory performance has been related to higher GMV in the prefrontal region (Goghari et al., 2014; Takeuchi et al., 2017) and larger surface area in superior and medial-orbital frontal gyri (Nissim et al., 2017), which lends support to our result that lower GMV in superior and middle frontal gyri relates to worse working memory performance in both adults and adolescents.

The identified top SNPs are SNPs in genes encoding long non-coding RNAs (lncRNAs) and SNPs in the genes MEF2C, CADPS2, CADM2, emphasizing potential roles of lncRNAs and a few biologically better interpretable genes in ADHD. The peak of the identified SNP component is coherent with the loci of the highest correlation with GMV in superior/middle frontal from the univariate analysis. They both highlight SNPs in a linkage disequilibrium structure in chromosome 5 located in lncRNA-encoding genes, including CTC-467M3.1, RP11_6N13.1, RP11_138J23.1, and RP11_38H17.1. RP11_6N13.1 has been associated with educational attainment (O’Connell et al., 2019) and broad depression (Howard et al., 2017), even though its biological function is understudied. Interestingly, the highlighted lncRNAs are highly expressed in testis and/or multiple brain regions, including the frontal cortex and anterior cingulate cortex based on the GTEx database (Ardlie et al., 2015). Even though it is unclear why these lncRNAs are expressed both in testis and brain, this is not a surprising finding given that brain and testis have demonstrated similar gene expression patterns by transcriptomic analyses in human (Guo et al., 2003), mouse (Guo et al., 2005), and zebrafish (Sreenivasan et al., 2008), suggesting that the proposed spICA can capture loci with modest effect size and similar expression patterns, and group them in one independent component.

Top SNPs identified in this work highlight MEF2C. Particularly, top SNP rs56144910 significantly regulates the methylation level of cg18498987 located in the 5’ untranslated region of MEF2C. MEF2C has gained increasing attention due to its role in neuronal development, function, and survival (Li et al., 2008a; Li et al., 2008b; Tang et al., 2005), and its involvement in neurodevelopmental disorders. In addition to its role in ADHD (Klein et al., 2020), MEF2C has been associated with autism spectrum disorder (Yingjun et al., 2017), intellectual disability (Savage et al., 2018; Sniekers et al., 2017), schizophrenia (Ripke et al., 2014) as well as Alzheimer disease (Lambert et al., 2013). Interestingly, Mef2c facilitated learning and memory in adult mice (Barbosa et al., 2008). After conditionally knocking out brain-specific Mef2c, mice were hyperactive and showed impaired motor coordination (Adachi et al., 2016). Importantly, MEF2C is highly expressed in the brain with the highest median expression in frontal cortex based on the GTEx database (Ardlie et al., 2015), which supports our result that the identified SNP component relates to superior and middle frontal cortex underlying working memory performance.

Another important gene emphasized by the identified top SNPs is CADM2. CADM2 has been associated with ADHD and executive function. Albayrak et al. identified that CADM2 related to hyperactivity/impulsivity in ADHD children (Albayrak et al., 2013). Importantly, a recent GWAS study (Ibrahim-Verbaas et al., 2016) on executive function and processing speed highlighted CADM2. Moreover, CADM2 is a known synaptic cell adhesion molecule that is vital for synapse organization. CADM2 is overexpressed in the brain with the highest expression in the frontal cortex (BA9), according to GTEx (Ardlie et al., 2015). Given that the frontal cortex has been well documented being involved in executive function (Funahashi and Andreau, 2013), our result, together with the literature, suggest that CADM2 is potentially modulating neural correlates in the frontal cortex of higher-level cognitive functions. Another critical gene highlighted by the identified top SNPs is CADPS2. CADPS2 is expressed in multiple tissues and widespread brain regions, including the occipital pole and frontal/temporal lobes (Cisternas et al., 2003). Moreover, genetic variants in CADPS2 have not only been associated with ADHD, but also with autism spectrum disorders and intellectual disability (Bonora et al., 2014), as well as Alzheimer’s disease (Velez et al., 2013).

The findings presented in this study should be considered in context with its strengths and limitations. This study used SNPs selected from the most recent ADHD GWAS study (Demontis et al., 2019), which provided us good candidates to investigate SNPs underlying brain alterations related to ADHD. However, samples from the NeuroIMAGE project included in this study were also utilized in the ADHD GWAS. It is worth noting that the ADHD GWAS summaries were only used to select SNPs and were not involved in the spICA analysis. Thus, the results identified by spICA are less likely to be inflated by overlapping samples. Another potential limitation is that spICA may overestimate the association between modalities. We have taken several steps to remedy and avoid this: 1) spICA monitors the entropy change introduced by the correlation optimization at each step and adaptively adjusts the step size for correlation optimization whenever the slope of the entropy is smaller than – 10^−5^; 2) we have used a replication dataset to validate the results from the discovery dataset.

In summary, we propose a new data fusion method, spICA, which imposes sparsity constraints implemented with the robust Hoyer projection and applied on components extracted within a pICA framework. The nonlinear Hoyer projection, together with the iterative reconstruction, offers access to a set of solutions otherwise inaccessible via traditional linear methods, like pICA. This results in sparser, less noisy sources than those obtained by linear transformation (Chi et al., 2013; Du et al., 2016; Le Floch et al., 2012; Vounou et al., 2010). On the other hand, incorrectly over enforcing sparsity may ruin meaningful signals. To alleviate this, we carefully preset Hoyer values for brain imaging and genetic data based on simulations with the desired number of signals, the expected SBR, and the total number of variables, assuming that the signal region forms a Laplacian distribution (to resemble the sparse nature of brain imaging/genetic sources) and the background forms a logistic distribution. We also conservatively preset step sizes (sMRI: 0.005, SNP: 0.0005) for sparsity enhancement (feasibility restoration) during optimization, and dynamically adjust them by monitoring the entropy change, thus prioritizing independence estimation. Simulation results demonstrated that spICA can robustly recover the designed sMRI-SNP associations, source maps, and loading matrices for both imaging and genetic modalities with improved accuracy compared to pICA. The accuracy improvement is more prominent for weak links, which make up most of the real-world associations between imaging phenotypes and genetics.

Applying the proposed spICA to brain imaging and genetic data of ADHD cohorts, we identified and replicated one SNP component related to GMV in superior and middle frontal gyri underlying working memory underperformance in adult and adolescent cohorts. The association was more significant in ADHD families than controls, and stronger in adults and older adolescents than younger ones. Top contributing SNPs reside in genes encoding lncRNA in chromosome 5 and a set of protein-coding genes (e.g., MEF2C, CADM2, CADPS2), which corroborate potential to modulate neuronal attributes underlying high-level cognition in ADHD.

## Acknowledgment

This study makes use of data from the Dutch NeuroIMAGE project, and the Dutch site of IMpACT (International Multi-center persistent ADHD Collaboration) project (IMpACT-NL). The NeuroIMAGE project was supported by NWO Large Investment grant 1750102007010 (Dr. Buitelaar), ZonMW Addiction: Risk Behaviour and Dependency Grant 60-60600-97-193 (Dr. Buitelaar), NWO Brain & Cognition: an Integrative Approach grant 433-09-242 (Dr. Buitelaar), NWO National Initiative Brain & Cognition 056-13-015 (Dr. Buitelaar), the EU FP7 grants TACTICS (278948), IMAGEMEND (602450), MATRICS (603016) and AGGRESSOTYPE (602805), EU IMI grant EU-AIMS (115300) and EU H2020 grant Eat2beNICE (728018) and grants from Radboudumc, University Medical Center Groningen, Accare, and VU University Amsterdam. The IMpACT-NL study acknowledges the following sources of support: The Netherlands Organization for Scientific Research (NWO), i.e., the NWO Brain & Cognition Excellence Program (grant 433-09-229) and the Innovation Program (Vici grant 016-130-669 to BF; Veni grant 016.196.115 to MH). Additional support was received from the European Community’s Seventh Framework Programme (FP7/2007 – 2013) under grant agreements n° 602805 (Aggressotype), n° 602450 (IMAGEMEND), and n° 278948 (TACTICS) as well as from the European Community’s Horizon 2020 Programme (H2020/2014 – 2020) under grant agreements n° 643051 (MiND), n° 728018 (Eat2beNICE) and n° 667302 (CoCA). The work was also supported by grants for the ENIGMA Consortium (grant number U54 EB020403) from the BD2K Initiative of a cross-NIH partnership and by the ECNP Network ADHD across the Lifespan. The authors also thank all participants involved in these two projects.

## Code Availability

The spICA code will be incorporated into the fusion ICA toolbox and can be freely download at https://trendscenter.org/software/.

