## supplemental materials for "Sparse parallel independent component analysis and its application to identify linked genomic and gray matter alterations underlying working memory impairment in attention-deficit/hyperactivity disorder"

---

**spICA algorithm pseudocode**

---

Require: datasets $\mathbf{X}_1$, $\mathbf{X}_2$, maximum number of the iteration T

Initialize the resularizer $\lambda$; preset Hoyer thresholds $q_1$, $q_2$; Hoyer sparsity enhancement step size $\Delta_{h1}$, $\Delta_{h2}$; unmixing matrices $\mathbf{W}_1$, $\mathbf{W}_2$; modality stopping flags $f_1$= FALSE, $f_2$= FALSE; entropy drop flags introduced by correlation optimization: $f_{c1}$= FALSE, $f_{c2}$= FALSE; $t_1 = 1, t_2 = 1$.

While ($t_1 < T$ & $f_1$ = FALSE) or ($t_2 < T$ & $f_2$ = FALSE) do

If $t_1 < T$ & $f_1$ = FALSE --Infomax on modality 1

Do an infomax update on $\mathbf{W_1}$ using stochastic gradient descent

If $\left\| \mathbf{W}_{1,\mathbf{t}_1} - \mathbf{W}_{1,\mathbf{t}_1-\mathbf{1}} \right\|_2^2 < \mathbf{10^{-6}}$

$f_1$ = TRUE

If the slope of the entropy of this modality is smaller than $-10^{-5}$

$f_{c1}$ = TRUE

If $\mathbf{t}_2 < \mathbf{T}$ **&** $f_2$ = FALSE --Infomax on modality 2

Do an infomax update on $\mathbf{W_2}$ using stochastic gradient descent

If $\left\| \mathbf{W}_{2,\mathbf{t}_2} - \mathbf{W}_{2,\mathbf{t}_2-\mathbf{1}} \right\|_2^2 < \mathbf{10^{-6}}$

$f_2$ = TRUE

If the slope of the entropy of this modality is smaller than $-10^{-5}$

$f_{c2}$ = TRUE

If $f_{c1}$ = TRUE or $f_{c2}$ = TRUE --Reduce the regularizer $\lambda$

$\lambda = 0.9 \times \lambda$

Compute the new source matrix $\mathbf{S}_1$ using the newly updated $\mathbf{W}_1$ as: $\mathbf{S}_1 = \mathbf{W}_1 \mathbf{X}_1$

If $f_1$ = FALSE --Sparsity optimization on modality 1

For each source (j-th row) $\mathbf{S}_{1,j}$ in $\mathbf{S}_1$ :

If the Hoyer index of the source $\mathbf{S}_{1,j}$ is less than the preset Hoyer value $q_1$:

Do Hoyer projection $\mathbf{S}_{H1,j} = \text{Hoyer}(\mathbf{S}_{1,j})$ with Hoyer step size $\Delta_{h1}$

Else: $\mathbf{S}_{H1,j} = \mathbf{S}_{1,j}$

If the slope of the entropy of this modality is smaller than $-10^{-5}$

    $\Delta_{h1} = 0.98 \times \Delta_{h1}$ --Reduce the Hoyer step size $\Delta_{h1}$

Reconstruct the data $\mathbf{X}_1$ using the cleaner source $\mathbf{S}_{H1}$: $\mathbf{X}_1 = \mathbf{W}_1^{+}\,\mathbf{S}_{H1}$

Compute the new source matrix $\mathbf{S}_2$ using the newly updated $\mathbf{W}_2$ as: $\mathbf{S}_2 = \mathbf{W}_2\,\mathbf{X}_2$

If $f_2$ = FALSE --Sparsity optimization on modality 2

    For each source (j-th row) $\mathbf{S}_{2,j}$ in $\mathbf{S}_2$

        If the Hoyer index of the source $\mathbf{S}_{2,j}$ is less than the preset Hoyer value $q_2$:

            Do Hoyer projection $\mathbf{S}_{H2,j} = \mathrm{Hoyer}(\mathbf{S}_{2,j})$ with Hoyer step size $\Delta_{h2}$

        Else: $\mathbf{S}_{H2,j} = \mathbf{S}_{2,j}$

If the slope of the entropy of this modality is smaller than $-10^{-5}$

    $\Delta_{h2} = 0.98 \times \Delta_{h2}$ --Reduce the Hoyer step size $\Delta_{h1}$

Reconstruct the data $\mathbf{X}_2$ using the cleaner source $\mathbf{S}_{H2}$: $\mathbf{X}_2 = \mathbf{W}_2^{+}\,\mathbf{S}_{H2}$

If $f_1$ = FALSE or $f_2$ = FALSE --Correlation optimization on modalities 1 and 2

    Update $\mathbf{W}_1$ and/or $\mathbf{W}_2$ based on correlation optimization with the regularizer $\lambda$

Increase the step $t_1$ by 1: $\mathbf{t}_1 = \mathbf{t}_1 + \mathbf{1}$

Increase the step $t_2$ by 1: $\mathbf{t}_2 = \mathbf{t}_2 + \mathbf{1}$

---

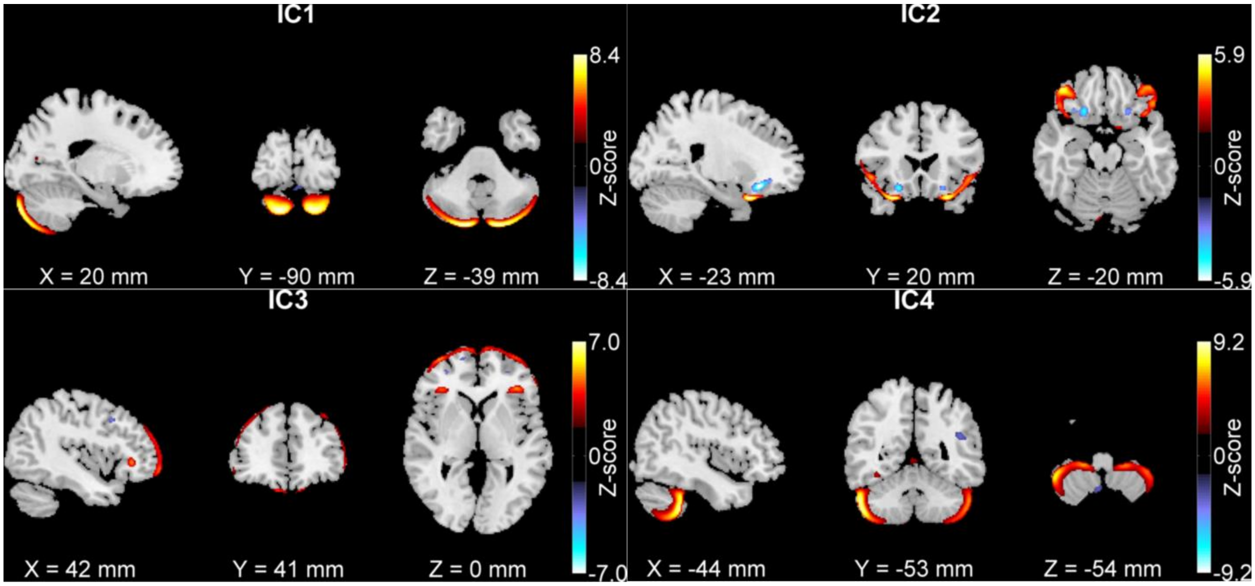


**Fig. S1.** Four GM components (ICs) significantly associated with either working memory or inattention symptoms (|Z| > 2.5). ICs 2-4 consistently associated with working memory performance/inattention in both ADHD adults and adolescents (Liu et al., 2020). Figure comes from the paper (Duan et al., 2018).

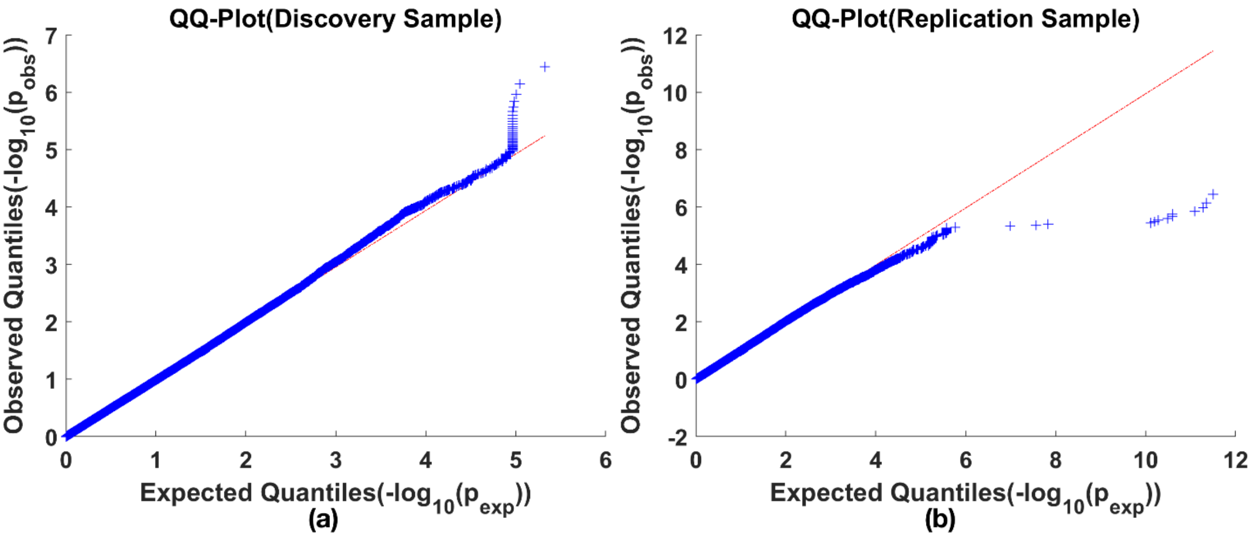


**Fig. S2**. Q-Q plots of sorted -log$_{10}$($p_{obs}$) values (ascending order, $p_{obs}$ values were obtained from univariate case vs. control MAF difference test) against sorted -log$_{10}$($p_{exp}$) values (ascending order, $p_{exp}$ were sampled from a uniform distribution) for (a) discovery and (b) replication samples.

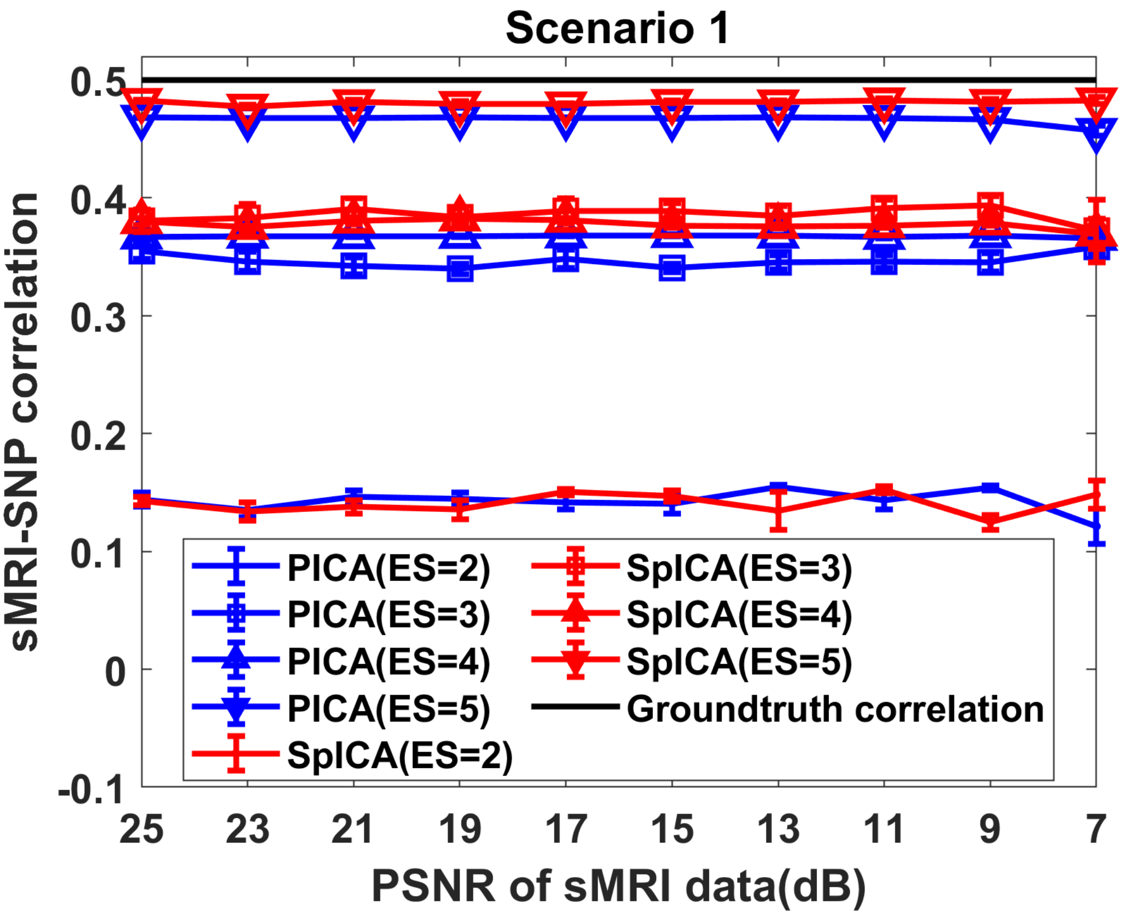


**Fig. S3**. Scenario 1: The sMRI-SNP correlations detected by spICA (red) and pICA (blue) while varying noise levels of sMRI components and effect size of SNP data. The ground-truth of the correlation was 0.5 (black).

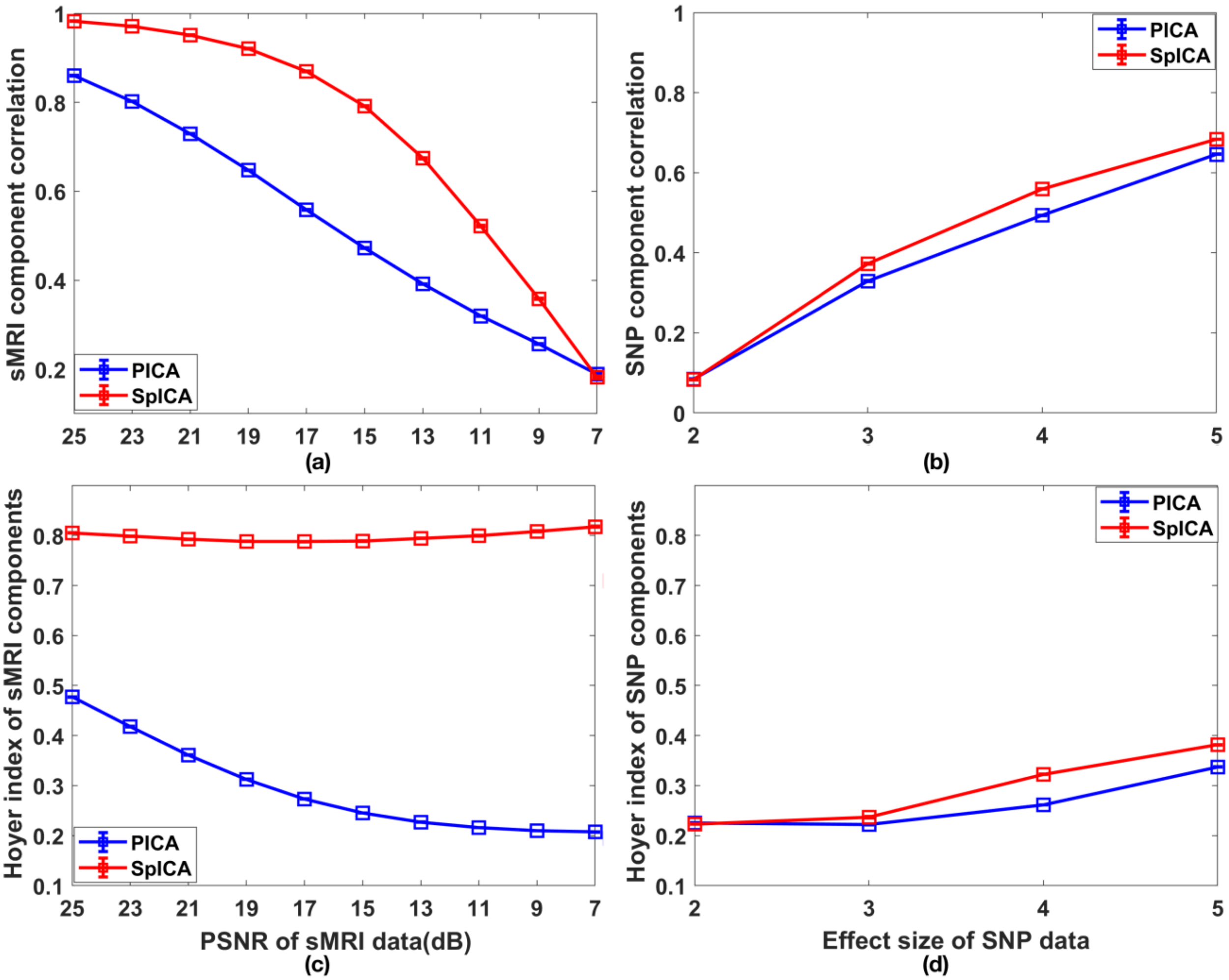


**Fig. S4**. Performance of spICA and pICA while varying PSNR values of white Gaussian noise superimposed on sMRI components and effect size of SNP data:(a) accuracy and (c) sparsity of recovered sMRI components; (b) accuracy and (d) sparsity of recovered SNP components (the designed sMRI-SNP association was fixed at 0.5).

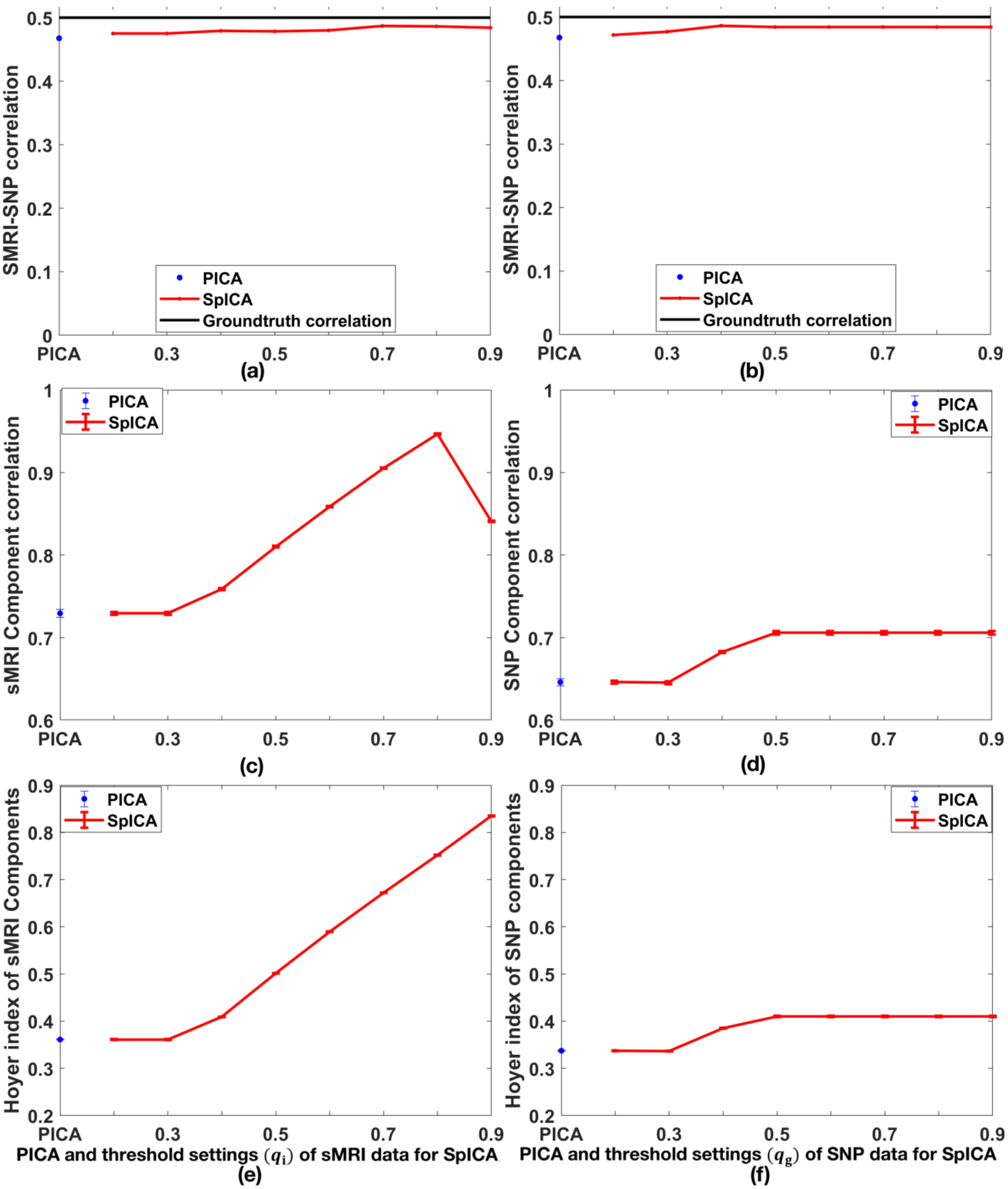


**Fig. S5**. Scenario 2: sMRI-SNP association detections from pICA and spICA with PSNR of 21dB and effect size of 5, varying Hoyer index thresholds from 0.2 to 0.9 for (a) sMRI data (varying $q_{\mathrm{i}}$ with fixed $q_{\mathrm{g}} = 0.4$, the ground-truth value for SNP data), (b) SNP data (varying $q_{\mathrm{g}}$ with fixed $q_{\mathrm{i}} = 0.85$, the ground-truth value for sMRI data); corresponding spICA and pICA performances including (c) accuracy and (e) sparsity of recovered sMRI components while varying $q_{\mathrm{i}}$ from 0.2 to 0.9; corresponding spICA and pICA performances including (d) accuracy and (f) sparsity of recovered SNP components while varying $q_g$ from 0.2 to 0.9 (the designed sMRI-SNP association was fixed at 0.5).

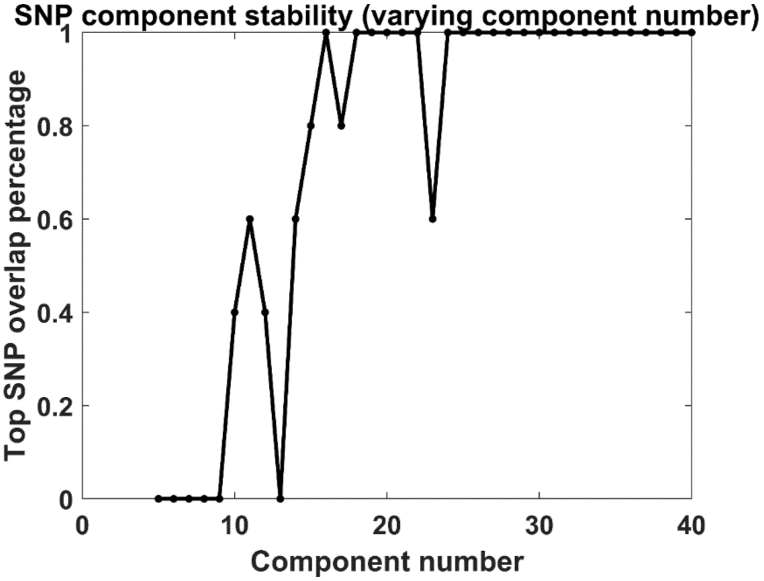


**Fig. S6**. Top SNP overlap ratio when SNP component number varied from 5 to 40.

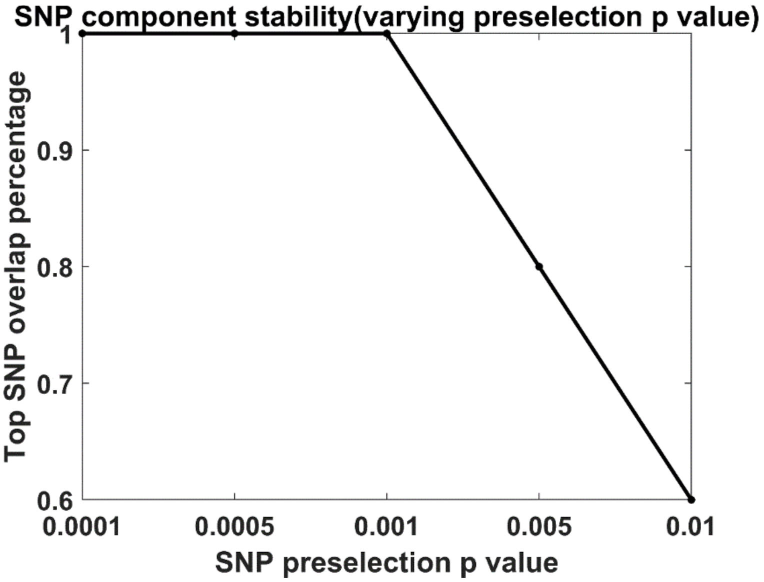


**Fig. S7**. Top SNP overlap ratio when SNP preselection p-value varied from 0.0001 to 0.01.

**TABLE S1.** Demographics of the adolescent population

| Variables | Diagnosis group (#) | | |
|---|---|---|---|
| | Healthy control (144) | Unaffected sibling (129) | ADHD (188) |
| Age | 14.50± 2.17 | 14.92±1.97 | 14.74± 2.32 |
| Sex(F/M) | 59/85 | 74/55 | 69/119 |
| Estimated IQ | 104.69± 13.54 | 99.22± 13.94 | 92.70± 16.09 |
| Inattention (IA) | 0.78± 1.71 | 1.35± 1.98 | 7.37±1.69 |
| Hyperactivity /Impulsiveness (HI) | 0.38± 1.14 | 0.97± 1.55 | 5.87±2.44 |
| Forward digit span (FW) | 8.78± 1.70 | 8.63± 1.63 | 7.84± 1.76 |
| Backward digit span (BW) | 6.07± 1.97 | 6.01± 1.72 | 4.95± 1.67 |
| History of stimulants | 1 | 11 | 106 |
| Scan site 1 | 101 | 71 | 88 |
| Scan site 2 | 43 | 58 | 100 |

**Table S2**, Top SNPs (|z|>2) in the identified SNP component, including their rsID, Chromosome, Base Pair Position, Gene Annotation, minor allele, z values.

| rsID | chromosome | bp position | gene annotation | minor allele | z value |
|---|---|---|---|---|---|
| rs786250 | 2 | 145701992 | TEX41 | T | 2.108993 |
| rs13006105 | 2 | 215205172 | SPAG16 | G | 2.318067 |
| rs10932536 | 2 | 215369762 | VWC2L | C | 3.989117 |
| rs10188314 | 2 | 215402926 | VWC2L | T | 4.389284 |
| rs4547674 | 3 | 43477118 | ANO10 | T | 2.39183 |
| rs737516 | 3 | 43558085 | ANO10 | T | 2.52676 |
| rs11921010 | 3 | 85434260 | CADM2 | T | 2.449831 |
| rs62250713 | 3 | 85513793 | CADM2 | A | 2.591646 |
| rs10433500 | 3 | 85546798 | CADM2 | G | 2.068441 |
| rs11928368 | 3 | 85661265 | CADM2 | T | -2.31886 |
| rs7690595 | 4 | 62464414 | ADGRL3 | G | 2.106978 |
| rs373098 | 5 | 88007261 | MEF2C-AS2 | C | -2.14915 |
| rs254778 | 5 | 88009593 | MEF2C-AS2 | G | -2.39045 |
| rs412458 | 5 | 88029627 | MEF2C | C | -2.35049 |
| rs664366 | 5 | 88055761 | MEF2C | A | -2.27184 |
| rs159950 | 5 | 88116319 | MEF2C | T | -2.00407 |
| rs2362108 | 5 | 88147771 | MEF2C | A | 2.798478 |
| rs61104616 | 5 | 88163771 | MEF2C | G | -2.1077 |
| rs3850651 | 5 | 88181109 | MEF2C | G | -2.60409 |
| rs2067663 | 5 | 88191635 | MEF2C | T | -2.60474 |
| rs56144910 | 5 | 88198557 | MEF2C | A | -2.36982 |
| rs1116527 | 6 | 149302053 | UST | T | 2.329897 |
| rs10231029 | 7 | 7907820 | UMAD1 | G | 3.751664 |
| rs6969177 | 7 | 7908235 | UMAD1 | G | 3.643153 |

| | | | | | |
|---|---|---|---|---|---|
| rs13438161 | 7 | 7914685 | UMAD1 | A | -2.05755 |
| rs11777844 | 8 | 1144023 | DLGAP2 | T | -2.32938 |
| rs935832 | 8 | 1145844 | DLGAP2 | T | 2.042755 |
| rs17669107 | 8 | 1152849 | DLGAP2 | C | 2.068176 |
| rs2811908 | 9 | 86247287 | IDNK | G | -2.00856 |
| rs7046039 | 9 | 112236879 | PTPN3 | T | 2.060341 |
| rs17292645 | 9 | 119503354 | ASTN2 | G | -2.24913 |
| rs6478248 | 9 | 119514613 | ASTN2 | A | -2.11502 |
| rs10821944 | 10 | 63785089 | ARID5B | G | 2.762281 |
| rs7902146 | 10 | 63801030 | ARID5B | C | 2.490487 |
| rs10821945 | 10 | 63801722 | ARID5B | G | -2.69097 |
| rs4948496 | 10 | 63805617 | ARID5B | C | 4.86717 |
| rs10761602 | 10 | 63813802 | ARID5B | G | 4.831801 |
| rs725529 | 10 | 63814070 | ARID5B | T | 4.347902 |
| rs11200360 | 10 | 123794424 | TACC2 | C | 3.129378 |
| rs66831679 | 12 | 73558582 | LINC02444 | T | -2.26856 |
| rs4297529 | 12 | 73569349 | LINC02444 | T | -2.25387 |
| rs55934137 | 12 | 73572753 | LINC02444 | G | -2.56101 |
| rs56150441 | 12 | 73583466 | LINC02444 | G | -3.04733 |
| rs7153641 | 14 | 91436255 | RPS6KA5 | T | -2.25177 |
| rs1286138 | 14 | 91485445 | RPS6KA5 | T | 2.371599 |
| rs8022176 | 14 | 91495427 | RPS6KA5 | C | 2.150126 |
| rs11847557 | 14 | 91507483 | RPS6KA5 | C | 3.396424 |
| rs1285992 | 14 | 91520438 | RPS6KA5 | A | 2.408921 |
| rs590247 | 15 | 47647560 | SEMA6D | C | 2.102224 |
| rs6047270 | 20 | 21122212 | KIZ | T | 2.042153 |
| rs2842198 | 1 | 43930738 | HYI | A | 2.193882 |
| rs10749820 | 1 | 73812598 | LINC01360 | C | 3.503534 |
| rs10789373 | 1 | 73965869 | | G | 3.199727 |
| rs10204857 | 2 | 146154475 | | G | 2.045052 |
| rs197261 | 2 | 161904823 | | G | 2.029682 |
| rs10804300 | 2 | 222883594 | | A | -2.22193 |
| rs4686038 | 3 | 6323162 | | G | 2.048103 |
| rs9859908 | 3 | 20489086 | | C | 2.082055 |
| rs6557063 | 5 | 92493318 | | C | -2.69197 |
| rs4703281 | 5 | 103397652 | | T | 5.868234 |
| rs1420864 | 5 | 103417386 | | C | 6.404969 |
| rs1420865 | 5 | 103419092 | | A | 6.81104 |
| rs12522020 | 5 | 103435822 | | T | 6.238597 |
| rs1882564 | 5 | 103462415 | | G | 6.549648 |
| rs12152847 | 5 | 103464999 | | C | 6.813598 |
| rs35090102 | 5 | 103480611 | | C | 5.71763 |
| rs13166522 | 5 | 103817315 | | A | 2.779196 |
| rs4235642 | 5 | 103818412 | | G | 2.534196 |
| rs6874138 | 5 | 103899596 | | T | 3.650389 |
| rs11738197 | 5 | 103913264 | | A | 3.604877 |
| rs325485 | 5 | 103995368 | | A | 3.870526 |
| rs325502 | 5 | 104008133 | | G | 3.814199 |
| rs12055234 | 5 | 104037760 | | A | 3.037056 |

| | | | | | |
|---|---|---|---|---|---|
| rs2400169 | 5 | 144526632 | | T | 2.379545 |
| rs2686805 | 7 | 68764352 | | A | -2.10463 |
| rs28522207 | 7 | 68764685 | | T | -2.39012 |
| rs28452470 | 7 | 121957582 | CADPS2 | A | -2.23288 |
| rs1475546 | 9 | 120533422 | | T | 2.093818 |
| rs997230 | 12 | 58748272 | | A | -2.6319 |
| rs66796775 | 12 | 73644444 | | A | -2.90352 |
| rs7963521 | 12 | 73828956 | | C | -2.46246 |
| rs17045844 | 12 | 79231591 | | G | -2.36244 |
| rs10220032 | 13 | 47615753 | | T | 2.109152 |
| rs9562700 | 13 | 47620956 | | A | 2.571834 |
| rs6561346 | 13 | 47621576 | | T | 3.288672 |
| rs7989860 | 13 | 47643335 | | G | 3.583762 |
| rs1313237 | 14 | 60848224 | | C | 2.245788 |
| rs34935520 | 14 | 61091401 | SALRNA1 | G | 3.156392 |
| rs10444728 | 14 | 63014025 | | G | 2.443636 |
| rs1959848 | 14 | 63045670 | | C | 2.309152 |
| rs17100152 | 14 | 63094740 | | C | 2.025206 |
| rs7201512 | 16 | 17913839 | | G | -2.04518 |
| rs6035821 | 20 | 21243293 | KIZ | T | 2.119121 |

**Table S3**, Summary of top SNPs regulation effects on gene expression ($p<10^{-3}$), including SNP ID and regulated genes as well as p values of the regulation effect obtained from (Ramasamy et al., 2014) .

| SNP ID | Gene name | p |
|---|---|---|
| rs17100152 | **RHOJ** | $4.00\times10^{-5}$ |
| rs2842198 | TIE1 | $3.60\times10^{-4}$ |
| rs2842198 | FLJ32224 | $5.10\times10^{-4}$ |
| rs10444728 | **RHOJ** | $5.10\times10^{-4}$ |
| rs1420864 | **NUDT12** | $7.90\times10^{-4}$ |
| rs1959848 | **RHOJ** | $8.00\times10^{-4}$ |
| rs13006105 | SPAG16, VWC2L | $8.50\times10^{-4}$ |
| rs11777844 | ERICH1, FLJ00290 | $8.80\times10^{-4}$ |
| rs12152847 | **NUDT12** | $9.20\times10^{-4}$ |
| rs2811908 | FRMD3 | $9.40\times10^{-4}$ |
| rs6478248 | ASTN2 | $9.60\times10^{-4}$ |

Note, top SNPs' regulation effect on expression of NUDT12 ($p$ = $1.32\times10^{-2}$, beta > 0) was also observed in the eQTL database for frontal cortex (Brodmann BA 9 region) in GTEx (Ardlie et al., 2015).

**Table S4**, Summary of 10 top SNPs regulation effects on methylation, with rsID, chromosome number, base pair location, minor allele and effect on GMV (local data), as well as the regulated CPG site and regulation effect on DNA methylation (DNAm) listed.

| SNP ID | Chr | BP position | Local SNP and GMV data | | Jaffe et al. (mQTL summary) | | |
|---|---|---|---|---|---|---|---|
| | | | Ref allele | Effect on GMV | CPG site | Annotation | Effect on DNAm |
| rs935832 | 8 | 1145844 | T | higher GMV | cg26999501 | | lower DNAm |
| rs17669107 | 8 | 1152849 | C | higher GMV | cg26999501 | | lower DNAm |
| rs1116527 | 6 | 149302053 | T | higher GMV | cg01163931 | UST | lower DNAm |
| rs2400169 | 5 | 144526632 | T | higher GMV | cg00055771 | | higher DNAm |
| rs7902146 | 10 | 63801030 | C | higher GMV | cg06318796 | ARID5B | lower DNAm |
| rs6557063 | 5 | 92493318 | C | lower GMV | cg06025774 | | lower DNAm |
| rs56144910 | 5 | 88198557 | A | lower GMV | cg18498987 | MEF2C | higher DNAm |
| rs11777844 | 8 | 1144023 | T | lower GMV | cg07699771 | | lower DNAm |
| rs28452470 | 7 | 121957582 | A | lower GMV | cg12076551 | | higher DNAm |
| rs2811908 | 9 | 86247287 | G | lower GMV | cg01266338 | C9orf103 | higher DNAm |

### S1, Genetic data preprocessing:

DNA extracted from blood was genotyped with the Illumina Psych Array for both NeuroIMAGE and IMpACT-NL projects. Pre-imputation quality control (QC) was performed to remove gender-mismatched samples and SNPs with minor allele frequency (MAF) < 0.01, call rate < 95%, and Hardy Weinberg Equilibrium < $1\times10^{-6}$. Imputation was performed based on ENIGMA protocol with 1000 genome as the reference. Only SNPs with imputation $r^2 > 0.3$ were included. We further removed participants with missing rate > 15% and SNPs with missing rate > 10% (yielding to 5,674,622 SNPs). Univariate case vs. control analysis was then performed; the obtained p values largely formed a uniform distribution (corresponding QQ plot is shown in Figure S2). Samples included in this study fell into a homogenous group (i.e., European ancestry). And we controlled for subgroup differences by using five genomic ancestry components.

### S2, Additional examination on spICA results:

#### *1, varying the number of SNP component from 5 to 40*

The identified SNP component in Figure 6(c) consistently showed up when SNP component number varied from 24 to 40 (the correlation coefficient between the identified SNP component and the matched one from different component order was between 0.60 and 0.67). Top 5 SNPs in the identified SNP component were still the top 5 in the matched component when SNP component order varied from 24 to 40. Figure S6 plotted the overlap ratio of the top 5 SNPs when SNP component number varied from 5 to 40. When component number is between 5 and 24, the variance of identified SNP component (Figure 6(c)) may not be included due to dimension deduction, thus resulting in low detection accuracy.

*2, varying SNP preselection p-value from 0.0001 to 0.01*

According to Chen's consistency measure, the SNP component numbers were estimated as 60, 37, 20, 50, 18 for SNP data with preselection p value threshold of 0.0001, 0.0005, 0.001, 0.005, 0.01, respectively. When SNP data preselection p-value varied from 0.0005 to 0.005, the identified SNP component consistently presented (the correlation between the identified SNP component and the matched one was between 0.53 and 0.67), and top 5 SNPs in the identified SNP component largely held as top 5 in the matched SNP component (Figure S7). When $p$ = 0.01, the correlation between the identified SNP component and the matched one is 0.44, and 3 out of 5 top SNPs were still top SNPs, which may be caused by the fact that too much SNPs were included when preselection p-value was 0.01 and the small variance in the identified SNP component may be dropped after dimension reduction.

*3, Applying heavy pruning ($r^2$=0.2) to SNP data*

When applying heavy pruning ($r^2$=0.2) to the SNP data with preselection $p < 1\times10^{-3}$, one GMV component significantly associated with one SNP component ($p = 5.91\times10^{-12}$, $\eta_p^2 = 0.19$), where the GMV component was highly correlated with GMV IC 1 in Figure 6(a) ($r$ = 1), and the SNP component had a correlation coefficient of 0.63 with the SNP component in Figure 6(c), and 4 out of 10 top SNPs were still presented as top SNPs in the matched one. New 6 top SNPs may result from the fact that p-value informed clumping in Plink software retained SNPs showing more significant case vs. control difference instead of those with larger weights in the SNP component. Noteworthily, the identified GMV-SNP component pair from $p < 1\times10^{-3}$, $r^2 < 0.2$ set was nominal significant in 317 subjects from ADHD families in adolescents ($p = 3.76\times10^{-2}$, $\eta_p^2 = 0.09$), indicating that the discovered GMV IC 1-SNP association was less likely biased by LD structure.

**S3, Brain regions of GMV IC 2:**

Figure 6(b) displayed the identified GMV IC 2 (z-scored), emphasizing inferior frontal gyrus ($|z| > 2.5$). For loadings of GMV IC 2, no significant GMV difference between ADHD patients and controls were observed in adults, and we observed significant GM reduction in inferior fontal gyri (hot colored region) for cases in adolescents with controlling for medication ($p = 9.59\times10^{-5}$, $t$ = 3.95, DF = 329). Loadings of GMV IC 2 showed no significant associations with working memory performances or symptom scores in 341 independent adults (it associated with forward and backward digit span task performances in a larger sample of 580 adults), but demonstrated significant and positive relation with forward digit span performance in adolescents ($p = 2.13\times10^{-2}$, $\eta_p^2 = 0.16$, beta > 0).

Ardlie, K.G., DeLuca, D.S., Segre, A.V., Sullivan, T.J., Young, T.R., Gelfand, E.T., Trowbridge, C.A., Maller, J.B., Tukiainen, T., Lek, M., Ward, L.D., Kheradpour, P., Iriarte, B., Meng, Y., Palmer, C.D., Esko, T., Winckler, W., Hirschhorn, J.N., Kellis, M., MacArthur, D.G., Getz, G., Shabalin, A.A., Li, G., Zhou, Y.H., Nobel, A.B., Rusyn, I., Wright, F.A., Lappalainen, T., Ferreira, P.G., Ongen, H., Rivas, M.A., Battle, A., Mostafavi, S., Monlong, J., Sammeth, M., Mele, M., Reverter, F., Goldmann, J.M., Koller, D., Guigo, R.,

McCarthy, M.I., Dermitzakis, E.T., Gamazon, E.R., Im, H.K., Konkashbaev, A., Nicolae, D.L., Cox, N.J., Flutre, T., Wen, X.Q., Stephens, M., Pritchard, J.K., Tu, Z.D., Zhang, B., Huang, T., Long, Q., Lin, L., Yang, J.L., Zhu, J., Liu, J., Brown, A., Mestichelli, B., Tidwell, D., Lo, E., Salvatore, M., Shad, S., Thomas, J.A., Lonsdale, J.T., Moser, M.T., Gillard, B.M., Karasik, E., Ramsey, K., Choi, C., Foster, B.A., Syron, J., Fleming, J., Magazine, H., Hasz, R., Walters, G.D., Bridge, J.P., Miklos, M., Sullivan, S., Barker, L.K., Traino, H.M., Mosavel, M., Siminoff, L.A., Valley, D.R., Rohrer, D.C., Jewell, S.D., Branton, P.A., Sobin, L.H., Barcus, M., Qi, L.Q., McLean, J., Hariharan, P., Um, K.S., Wu, S.P., Tabor, D., Shive, C., Smith, A.M., Buia, S.A., Undale, A.H., Robinson, K.L., Roche, N., Valentino, K.M., Britton, A., Burges, R., Bradbury, D., Hambright, K.W., Seleski, J., Korzeniewski, G.E., Erickson, K., Marcus, Y., Tejada, J., Taherian, M., Lu, C.R., Basile, M., Mash, D.C., Volpi, S., Struewing, J.P., Temple, G.F., Boyer, J., Colantuoni, D., Little, R., Koester, S., Carithers, L.J., Moore, H.M., Guan, P., Compton, C., Sawyer, S.J., Demchok, J.P., Vaught, J.B., Rabiner, C.A., Lockhart, N.C., Ardlie, K.G., Getz, G., Wright, F.A., Kellis, M., Volpi, S., Dermitzakis, E.T., Consortium, G., 2015. The Genotype-Tissue Expression (GTEx) pilot analysis: Multitissue gene regulation in humans. Science 348, 648-660.

Duan, K., Chen, J., Calhoun, V.D., Lin, D., Jiang, W., Franke, B., Buitelaar, J.K., Hoogman, M., Arias-Vasquez, A., Turner, J.A., Liu, J., 2018. Neural correlates of cognitive function and symptoms in attention-deficit/hyperactivity disorder in adults. Neuroimage Clin 19, 374-383.

Liu, J., Duan, K., Jiang, W., Rootes-Murdy, K., Schoenmacker, G., Buitelaar, J.K., Hoogman, M., Oosterlaan, J., Hoekstra, P.J., Heslenfeld, D.J., Hartman, C.A., Calhoun, V.D., Arias-Vasquez, A., Turner, J.A., 2020. Gray matter networks associated with cognitive deficit in ADHD across adolescence and adulthood. medRxiv, 2020.2004.2022.20059808.

Ramasamy, A., Trabzuni, D., Guelfi, S., Varghese, V., Smith, C., Walker, R., De, T., Coin, L., de Silva, R., Cookson, M.R., Singleton, A.B., Hardy, J., Ryten, M., Weale, M.E., Consortium, U.B.E., Consor, N.A.B.E., 2014. Genetic variability in the regulation of gene expression in ten regions of the human brain. Nature Neuroscience 17, 1418-1428.
